# Structural insights into a single semi-clathrate hydrate formed in a confined environment of porous protein crystal

**DOI:** 10.1101/2023.06.21.546005

**Authors:** Basudev Maity, Jiaxin Tian, Tadaomi Furuta, Satoshi Abe, Takafumi Ueno

**Author notes:** Corresponding authors. (TF); (TU). Equal contribution (Basudev Maity, Jiaxin Tian).

## Abstract

Water behavior on protein surfaces influences protein structure and function. Antifreeze Proteins (AFPs) have been intensively studied in context of biological cytotechnology. AFPs inhibit growth of ice microcrystals by forming unique water-cluster networks which are influenced by protein surface morphology and hydrophobicity. Such unique water-cluster networks have been identified as semi-clathrate structures in crystals and are believed to be stabilized by intermolecular interactions within the confined environment. However, there is little atomic-level information about the process of formation of semi-clathrates and the structural units of water-clathrate networks. We identified a single semi-clathrate formed on the pore surface of ferritin crystal which has a structure similar to that of a natural AFP. Comparison of ferritin mutants and determination of temperature-dependent structures revealed that semi-clathrate water molecules on an ⍺-helix undergo structural alterations with increasing temperature. Lowering the temperature regenerates the semi-clathrate structure. Water molecules hydrogen-bonded to main chain carbonyl groups are stably immobilized at room temperature and serve as starting points for clathrate formation. These findings provide a mechanistic understanding of water networks in AFPs and guidelines for designing new cryomaterials.

## Introduction

The dynamics of water and hydrogen bonding networks among biomolecules such as proteins and nucleic acids are fundamental to life.(1–3) The properties of water networks are influenced by hydrophobicity, crowding, and confinement of biomolecules.(4–6) The presence of highly structured interfacial water molecules correlates with specific functions and structures of proteins.(7, 8) Understanding the behavior of proteins in response to perturbations that alter the structure and dynamics of water molecules associated with biomolecules is essential, and vice versa. Thus, it is important to understand the changes in water dynamics induced on protein surfaces. NMR, X-ray and neutron scattering, and molecular simulations have been used to study issues related to the structure and dynamics of water at protein interfaces.(1, 9–12) For example, studies on natural antifreeze proteins (AFPs) and their analogs are expected to drive development of applications in the cryopreservation of cells, tissues, organs, food preservation, and antifreeze materials.(13–15) Therefore, to understand the role of water in life at the atomic level, it is necessary to unambiguously determine structures and interactions at the protein-water interface and systematically analyze the interaction geometry between protein and water molecules, as well as the local and global distribution of water molecules on protein surfaces. Polygon structures of water clusters at protein interfaces are closely related to protein surface structures.(16–18) Crystal structures of proteins tend to have polygonal hydrogen bonding networks on their surfaces. For example, a series of pentagonal water clusters surrounding Leu-18 have been observed within the crystal of crambin, a small hydrophobic protein.(18) Formation of similar pentagonal clusters in the binding cavity of streptavidin has been identified in MD simulations.(19) In addition, water cluster structures have been found to be essential in AFPs. Sun et al. reported that a 33 kDa AFP known as “Maxi,” has a unique shell consisting of over 400 water molecules, forming semi-clathrates composed of pentagonal water clusters (Figure 1a).(20, 21) It is believed that these networks are responsible for the interactions between proteins and the ice crystal surface.(17, 22) However, the influence of local protein structure on semi-clathrate formation is not well understood because it is difficult to design and construct single semi-clathrates on protein surfaces and observe them at atomic-level resolution. Moreover, the factors that promote semi-clathrate formation on protein surfaces have been studied computationally.(23) Few semi-clathrate structures have been identified experimentally because they exist in narrow spaces or are anchored by interactions with many amino acid residues.(17, 18, 20, 24) Thus, the mechanisms of formation of semi-clathrate structures are not yet fully understood. Interactions of ferritin with ice nuclei have been studied by analytical methods such as dynamic light scattering and differential scanning calorimetry.(25, 26) However, the presence of a semi-clathrate structure on the surface of ferritin has not been previously reported.

**Figure 1.**
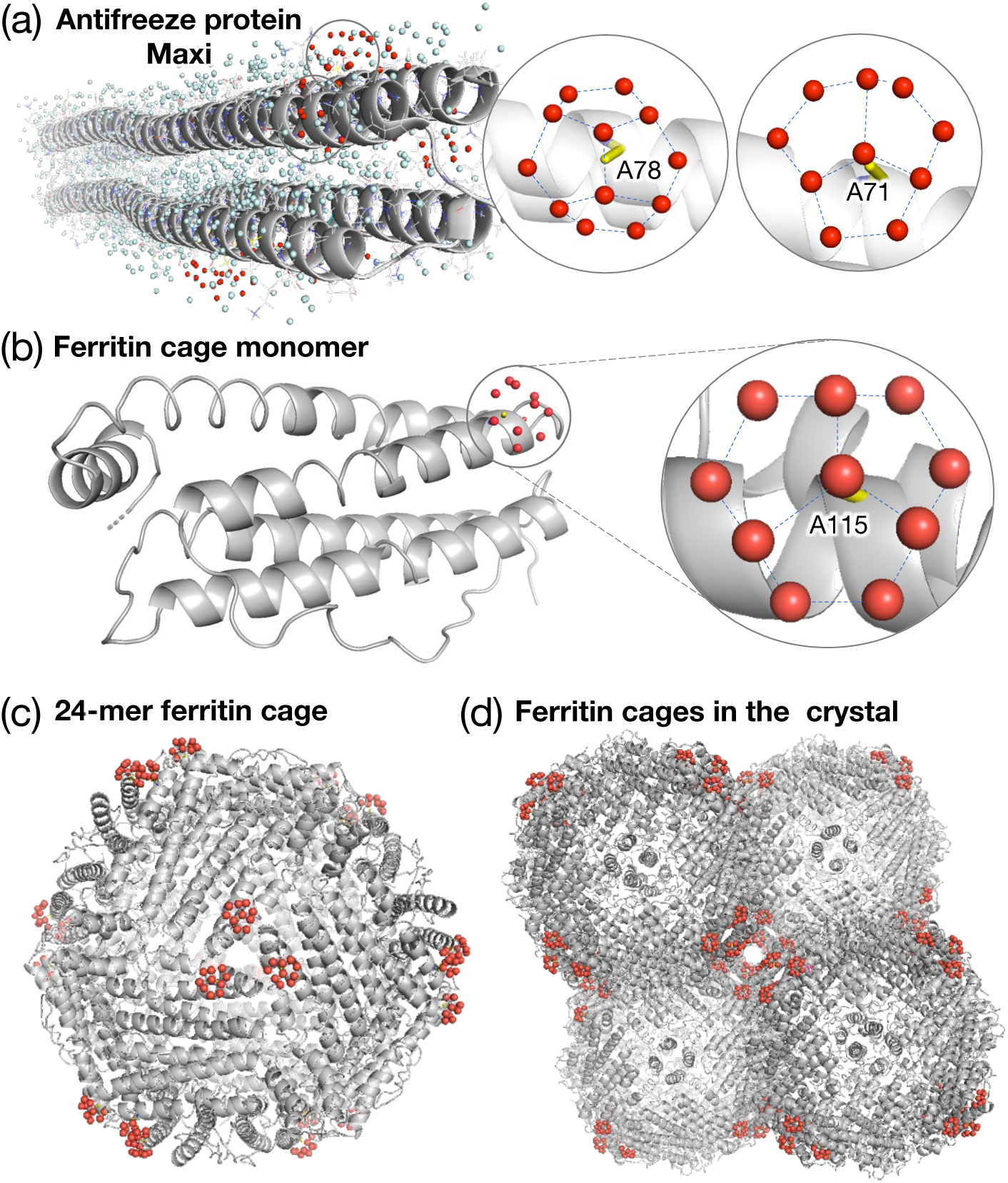
Semi-clathrate hydrates in proteins. (a) Structure of a hyperactive type-I antifreeze protein, Maxi (four-helix bundle) (PDB: 4KE2). Insets show two selected semi-clathrate hydrate structures at Ala71 and Ala78, where the semi-clathrate waters (red spheres) form fused pentagonal rings over Ala residues. (b) Structure of ferritin cage monomer (PDB: 8I6L). Inset shows the presence of a single semi-clathrate hydrate structure over Ala115 as in (a). (c) 24-mer ferritin cage structure. The semi-clathrates (red spheres) are located near the three-fold symmetric channels (outside). (d) Cross section of four ferritin cages. In the arrangement of ferritin cages in crystal, solvent channels and semi-clathrates are locate close to each other. Figures are prepared in pymol.

Herein, we describe the structural details of a single semi-clathrate in a crystal of ferritin which is formed over one specific alanine residue which is exposed to a large solvent channel (Figure 1) in an arrangement similar to the framework observed in Maxi. The semi-clathrate is hydrogen-bonded to the main chain carbonyls of an ⍺-helix and water molecules are tightly anchored within the crevice of the molecular interface. The influence of temperature on the clathrate structure indicates that some hydrogen bonds are maintained at temperatures close to room temperature. Moreover, replacement of alanine with other amino acid residues causes the semi-clathrates to be transformed into different polygonal clusters, with retention of the hydrogen bonds. This finding indicates that the hydrogen bonding network underlying the clathrate contributes significantly to its formation and stability. Through high-resolution crystallography, we have succeeded in constructing a model system that can be used to study the formation of water clusters on protein surfaces in detail. The crystal engineering of ferritin presented in this study is expected to enhance our understanding of the mechanisms of formation of semi-clathrates in antifreeze proteins and inspire molecular design of new cryomaterials.

## Results

### Structure of a single semi-clathrate hydrate in a wild-type ferritin crystal

Recombinant wild-type L-chain ferritin from horse spleen (FrWT) was prepared and crystallized in the presence of CdSO_4_ and (NH_4_)_2_SO_4_ (Figure S1a).(27, 28) The X-ray structure of FrWT was determined at 1.5 Å resolution at –180℃. The crystallographic data and refinement statistics are listed in Table S1. The overall 24-mer structure in this study (PDB: 8I6L) is similar to the previously reported structure (PDB: 1DAT)(29) with a root mean square deviation (RMSD) of 0.187 Å for the C^⍺^ of the main chain carbon atoms. We analyzed the water networks near the alanine residues in the ferritin cage (Figure S2) because a semi-clathrate structure was previously found to form over Ala in the natural antifreeze protein known as Maxi.(20) Out of a total of fifteen alanine residues present in the ferritin cage monomer, we observed the presence of a single semi-clathrate structure containing ten water molecules (W1∼W10) over Ala115 (Figure 2). This semi-clathrate structure is located near the exit of the three-fold symmetric channel on the outer surface of the ferritin cage (Figure 2a). The structure is formed of three fused pentagonal rings which are roughly planar with a deviation of dihedral angle below 20° to the plane containing three edge water molecules (Figure S3). The three successive edges within the pentagonal water ring form dihedral angles ranging from 116° to 124° (Figure S3). The distances of the semi-clathrate water molecules from the central Ala115 (C^β^) are in the range of 3.6-4.4 Å (Table S2). The average B factor of the 10 semi-clathrate water molecules is 24.8 Å^2^ (Table S3). The observed oxygen-oxygen distances in the semi-clathrate are in the range of 2.44-3.28 Å, which is within hydrogen-bonding range (Table S4).(30–32) The observed semi-clathrate is similar to the semi-clathrate of Maxi (Figure 1a and Table S4).(20) The semi-clathrate water molecules W1∼W10 at Ala115 interact with the surrounding side chains, main chains and anchoring water molecules which were assumed to stabilize the semi-clathrate structure (Figure 2 and Table 1). W1, located at the vertex of the three pentagonal water rings, is stabilized only by the neighboring semi-clathrate W2, W5 and W8 molecules (Figure 2a). W1-W2, W1-W5 and W1-W8 form the edges between the pentagons and these molecules are only stabilized by the surrounding semi-clathrate water molecules. Out of 10 semi-clathrate water molecules, only W9 directly forms a hydrogen bond with the side chain of Asp112 with a W9-O^ᵟ^(Asp112) distance of 2.73 Å (Table 1 and Figure S4). On the other hand, W4 and W7 directly form hydrogen bonds with the main chain carbonyl oxygen atoms of Ala115 and Asp112 with distances of 2.75 Å and 2.76 Å, respectively. In addition, we observed several anchoring water molecules which form hydrogen bonding networks between the semi-clathrate water molecules and surrounding residues (cyan spheres in Figure 2). W3 and W4 are stabilized by His114 and Ser118, respectively, by anchoring one water molecule each (Figure 2a and Table 1). Moreover, W6 is stabilized weakly by Gln120 by anchoring of two water molecules. Similarly, W10 is also stabilized by both Gln108 and Leu111 by anchoring two water molecules via a shared water molecule. These main interactions above are expected to provide stability to the semi-clathrate structure.

**Figure 2.**
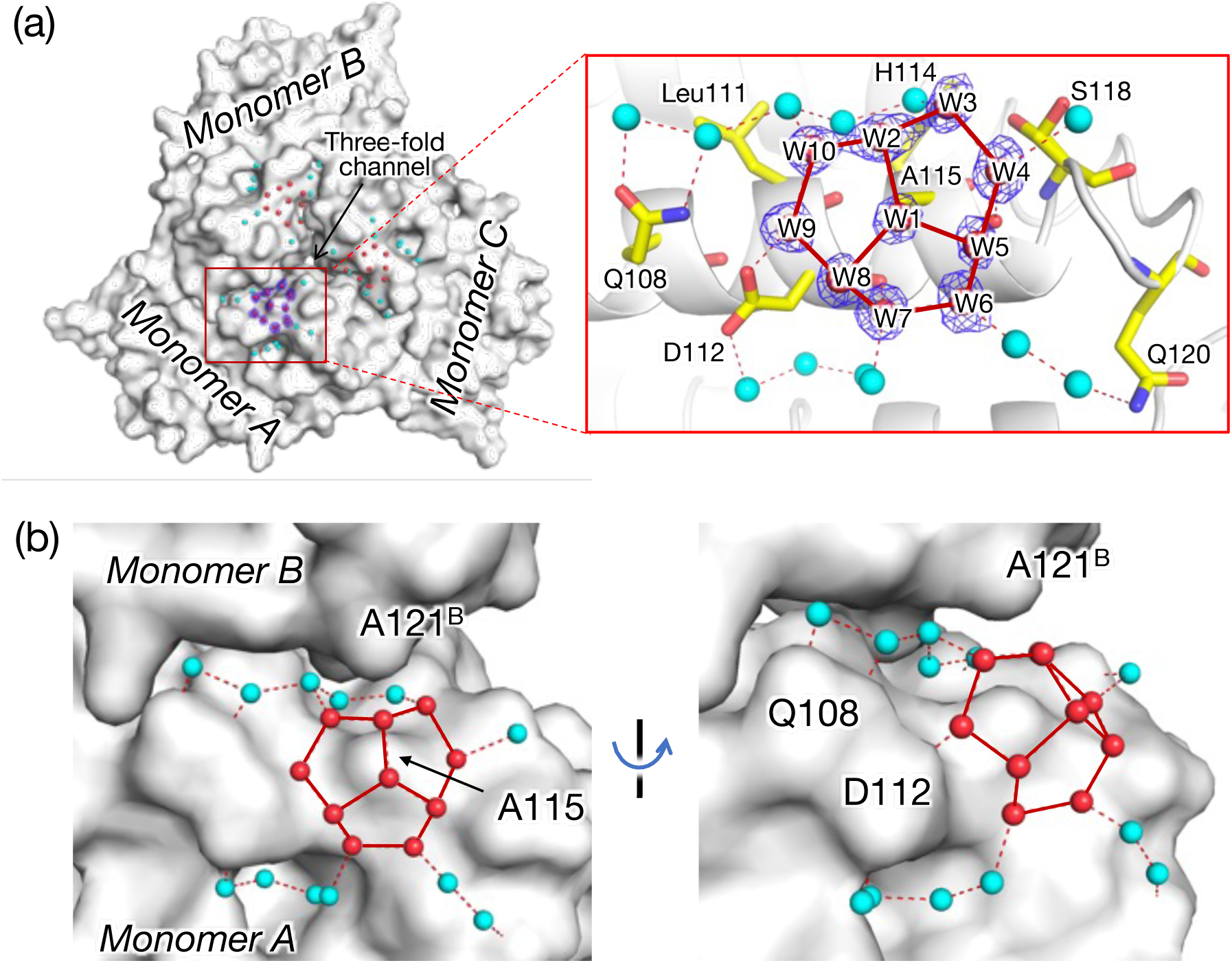
Detailed structure of the semi-clathrate water network. (a) Surface model of the ferritin trimer near the three-fold symmetric channel surface at -180℃. Inset shows the detail structure of the semi-clathrate and interactions with surrounding residues and water molecules. 2*F*_o_-*F*_c_ maps of the selected water molecules are shown in blue mesh and contoured at 1.0σ. (b) Semi-clathrate water network at the interface between adjacent monomers. The semi-clathrate and surrounding waters are depicted in red and cyan spheres, respectively. The solid and dotted lines represent hydrogen bonds in the semi-clathrate and others, respectively. Figures are prepared in pymol.

**Table 1.**
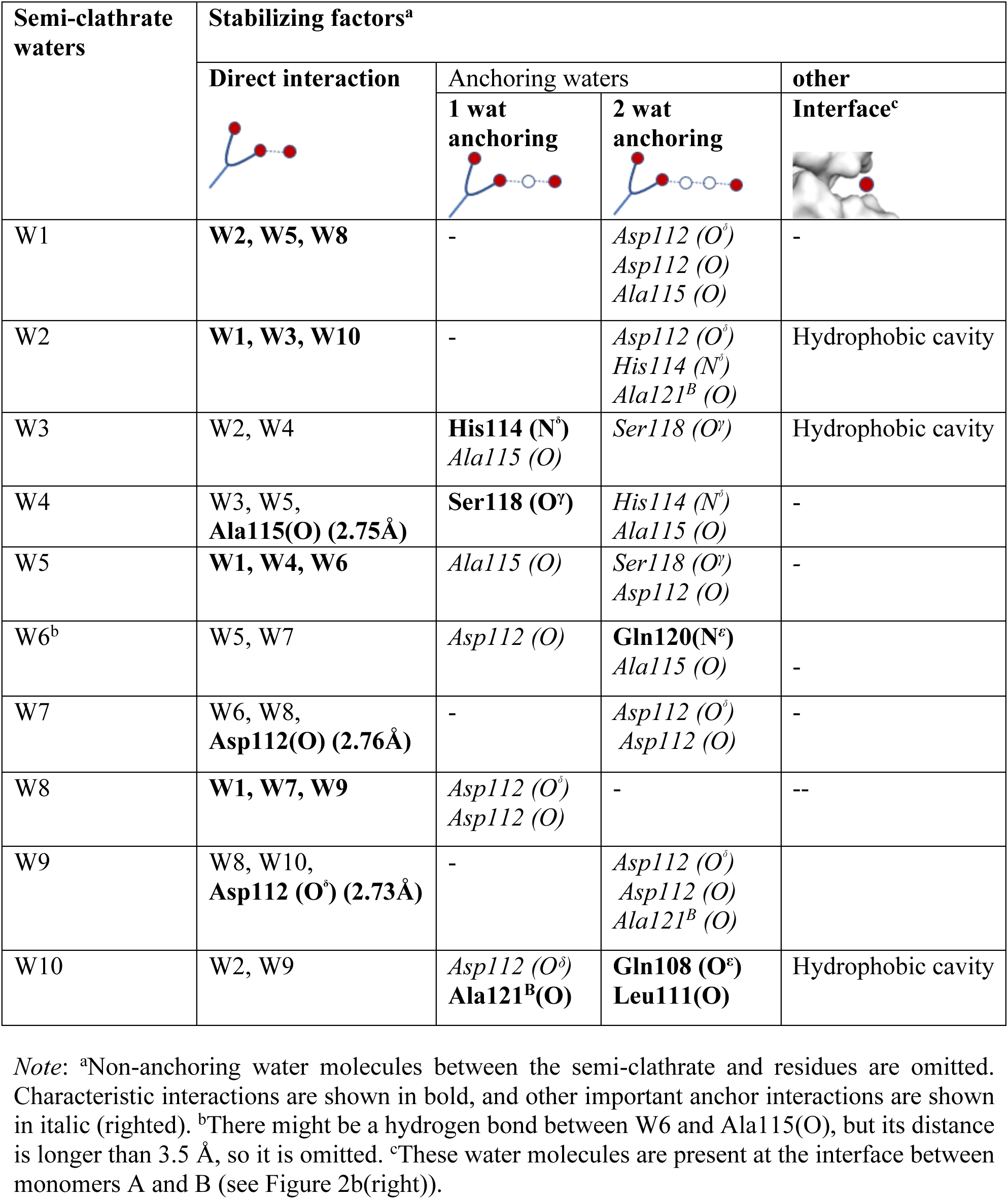
Stabilization factors of the semi-clathrate waters (W1-W10) in FrWT at -180°C.

| Semi-clathrate waters | Stabilizing factors<sup>a</sup> |  |  |  |
| --- | --- | --- | --- | --- |
|  | Direct interaction | Anchoring waters |  | other<br>Interface<sup>c</sup> |
|  |  | 1 wat anchoring | 2 wat anchoring |  |
| W1 | **W2, W5, W8** | - | *Asp112 (O<sup>δ</sup>)*<br>*Asp112 (O)*<br>*Ala115 (O)* | - |
| W2 | **W1, W3, W10** | - | *Asp112 (O<sup>δ</sup>)*<br>*His114 (N<sup>δ</sup>)*<br>*Ala121<sup>B</sup> (O)* | Hydrophobic cavity |
| W3 | W2, W4 | **His114 (N<sup>δ</sup>)**<br>*Ala115 (O)* | *Ser118 (O<sup>γ</sup>)* | Hydrophobic cavity |
| W4 | W3, W5,<br>**Ala115(O) (2.75Å)** | **Ser118 (O<sup>γ</sup>)** | *His114 (N<sup>δ</sup>)*<br>*Ala115 (O)* | - |
| W5 | **W1, W4, W6** | *Ala115 (O)* | *Ser118 (O<sup>γ</sup>)*<br>*Asp112 (O)* | - |
| W6<sup>b</sup> | W5, W7 | *Asp112 (O)* | **Gln120(N<sup>ε</sup>)**<br>*Ala115 (O)* | - |
| W7 | W6, W8,<br>**Asp112(O) (2.76Å)** | - | *Asp112 (O<sup>δ</sup>)*<br>*Asp112 (O)* | - |
| W8 | **W1, W7, W9** | *Asp112 (O<sup>δ</sup>)*<br>*Asp112 (O)* | - | -- |
| W9 | W8, W10,<br>**Asp112 (O<sup>δ</sup>) (2.73Å)** | - | *Asp112 (O<sup>δ</sup>)*<br>*Asp112 (O)*<br>*Ala121<sup>B</sup> (O)* |  |
| W10 | W2, W9 | *Asp112 (O<sup>δ</sup>)*<br>**Ala121<sup>B</sup>(O)** | **Gln108 (O<sup>ε</sup>)**<br>**Leu111(O)** | Hydrophobic cavity |

*Note:* <sup>a</sup>Non-anchoring water molecules between the semi-clathrate and residues are omitted. Characteristic interactions are shown in bold, and other important anchor interactions are shown in italic (righted). <sup>b</sup>There might be a hydrogen bond between W6 and Ala115(O), but its distance is longer than 3.5 Å, so it is omitted. <sup>c</sup>These water molecules are present at the interface between monomers A and B (see Figure 2b(right)).

Apart from such polar interactions with side chains, the main chain and anchoring water molecules that could be seen on the ferritin monomer, one side of the semi-clathrate structure which encompasses W2, W3 and W10 is located at the interface of two ferritin monomers (Figure 2b). The hydrophobic cavity generated by Ala115 (monomer A) and Ala121B (monomer B) stabilizes a unique six-membered water ring which has one water molecule located out-of-plane as a result of a hydrogen bond interaction (2.92 Å) with carbonyl oxygen atom of Ala121B (Figure 2b and S5). The observed Ala115 (C^β^) - Ala121B (C^β^) distance is 5.85 Å. The six-membered ring attached to the semi-clathrate between Ala115 and Ala121B is similar to the fourth pentagonal water ring located above Ala78 in Maxi (Figure 1a and S5).

### Roles of the central residue on the semi-clathrate formation

The semi-clathrate water structure described above is located above Ala115 in FrWT (Figure 2a). To explore the role of this Ala115 residue, we prepared a series of ferritin mutants (Figure 3a) with Ala115 replaced by Val, Thr and Gly (Fr-A115V, Fr-A115T and Fr-A115G, respectively). The mutants were purified and crystallized using the same methods used for FrWT (Figure S1b). The structures were refined with 1.5-1.6 Å resolution and the RMSDs calculated for the main chain C^⍺^ atoms of FrWT are 0.10-0.147 Å, suggesting that the general structures of the mutants are similar to the general structure of FrWT.

**Figure 3.**
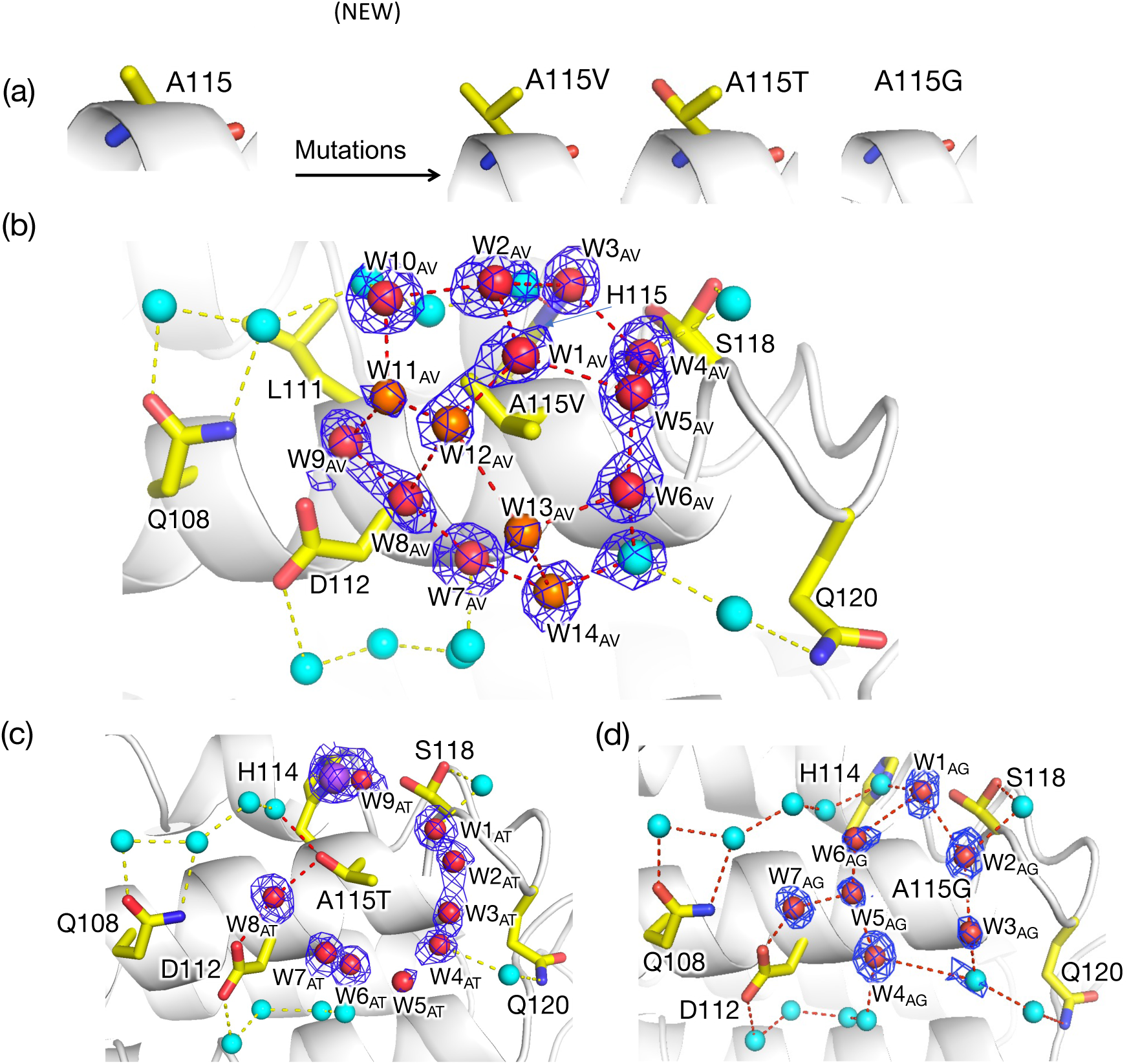
Changes in semi-clathrate water structures due to mutations at central Ala115. (a) Design scheme for mutations at residue 115. (b) Water networks over the central Val115 in Fr-A115V mutant. (c) Water networks over the central Thr in Fr-A115T mutant. Na^+^ ion (assigned due to its high electron density compared to water molecule) is depicted by purple sphere. (d) Water networks over the central Gly in Fr-A115G mutant. Red and cyan spheres represent water molecules near the central residue and surrounding water molecules, respectively, and orange spheres in (b) represent the additional ones appeared due to Val mutation. The blue mesh represents the 2*F*_o_-*F*_c_ maps at 1.0σ for the corresponding water molecules. Figures are prepared in pymol.

In the Fr-A115V mutant, we observed formation of a wider semi-clathrate structure above Val115 (Figure 3b). The ten water molecules observed in the semi-clathrate of FrWT are retained in the A115V mutant, but with significant deviations (red spheres in Figure 3b and S6). Four additional water molecules (W11_AV_, W12_AV_, W13_AV_ and W14_AV_) are located above V115 to form the semi-clathrate water structure, which consists of quadrangular rings (orange spheres in Figure 3b and S6) as well as pentagonal rings. The observed wat-wat distances (W1_AV_∼W14_AV_) are in the range of 2.35-3.48 Å (Table S5). The water network of anchoring water molecules interacting with the surrounding residues Gln108, Leu111, Asp112, His114, Ser118 and Gln120 remains in a position similar to that of FrWT (cyan spheres in Figure 3b and S6). The distance (5.05 Å) between W1 and the central A115V (C^β^) was found to be longer than that of WT-Fr (4.00 Å) (Table S2). This indicates that the water network in this mutant is formed farther from the protein surface due to the additional steric hinderance provided by the bulky side chain of valine at position 115 (Figure S6). Due to this shift, additional water molecules W11_AV_, W12_AV_, W13_AV_ and W14_AV_ participate in the semi-clathrate water network and are accompanied by large movements (from FrWT) of neighboring W1_AV_, W2_AV_, W5_AV_, W6_AV_ and W10_AV_, which cover the hydrophobic Val115 residue (Figure 3b and S6, and Table S2). In addition, due to the expansion of this water molecule network, several water molecules have higher B-factors (possible low occupancy) as well as shorter wat-wat distances, in comparison to the water molecules of the network of FrWT (Table S5). On the other hand, no significant change was observed at the interface side where His114 and Ser118 interact with the semi-clathrate through indirect hydrogen bonds. These results indicate that the semi-clathrate structure is formed over the hydrophobic residues. The semi-clathrate water molecules are positioned according to their distances from the protein surface and their hydrogen bonding, and are immobilized upon entering the crevice.

To investigate the role of hydrophobicity at the central core of semi-clathrate structure, we replaced the hydrophobic Val115 reside with Thr which is structurally similar but polar (the Thr residue has a methyl group and a hydroxyl group whereas Val has two methyl groups (Figure 3a)). In this Fr-A115T mutant, a semi-clathrate water structure was not observed above Thr115. This suggests that a central hydrophobic core is required to form the semi-clathrate water structure. Nine peripheral water molecules surrounding A115T are located at positions similar to the positions observed in the Fr-A115V mutant (Figure 3b and 3c), where several water molecules including W1_AV_, W11_AV_∼W13_AV_ in Fr-A115V are not observed. The OH group of the central Thr115 reside was found to form hydrogen bonds with W8_AT_, which also interacts with Asp112, giving W8_AT_- O*^v^*(Thr115) and W8_AT_-O^ᵟ^(Asp112) distances of 3.0 Å and 2.7 Å, respectively (Table S8). Thr115 also interacts with an additional water molecule at a distance of 3.2 Å. This additional water molecule is located at the interface of two ferritin monomers. The surrounding water networks involving Gln108, Leu111, Asp112, His114, Ser118, and Gln120 are essentially the same as those of the Fr-A115V mutant (Figure 3b and 3c). This observation suggests that the peripheral water networks are stabilized in the semi-clathrate in the Fr-A115V mutant, and that a hydrophobic central residue is necessary to form these water networks. Although Thr is structurally similar to Val, the presence of the hydroxyl group of Thr prevents formation of the semi-clathrate in the Fr-A115T mutant. Similarly, Thr residues are not covered with semi-clathrate water molecules in the natural antifreeze protein Maxi (pdb: 4KE2). Considering the coordination geometry and noting that the density of a water molecule coordinated by His114 appears larger than neighboring water molecules, we tentatively assign this observation as indicating the presence of a sodium ion (Figure 3c and Table S8). This interpretation agrees with the previously reported association between metal ions and water cluster structures.(24, 33) In addition, a six membered water ring at the interface of two monomers was not observed for the Fr-A115T mutant (Figure S7b). This also suggests that the central hydrophobic residue plays an important role in development of water networks at the monomer-monomer interface.

Moreover, we replaced the Ala115 with Gly to provide a minimal side chain. In this Fr-A115G mutant, the semi-clathrate water structure seen in FrWT and the Fr-A115V mutant is absent, but seven peripheral water molecules surrounding Gly115 were observed (Figure 3e). The observed wat-wat distances are in the range of 2.34-3.81 Å (Table S8). The anchoring water networks interacting with the surrounding residues Gln108, Leu111, Asp112, His114, Ser118, and Gln120 are also conserved in this mutant. Similar to the observations of the Fr-A115T mutant described above, a six membered water ring was not observed for the A115G mutant (Figure S7c). These results suggest that the formation of the semi-clathrate structure requires a centrally located hydrophobic residue of appropriate size.

### Roles of the surrounding residues on the semi-clathrate formation

As described above, the residues Asp112, His114, and Ser118, are significantly involved in formation of the semi-clathrate structure. Therefore, we designed three additional mutants, Fr-D112A, Fr-H114A, Fr-S118A, to investigate the effects of these residues. Each of these mutants were expressed and purified using the same techniques used to obtain FrWT. However, we could not obtain Fr-D112A as the 24-mer. The other mutants were characterized by native-PAGE, SDS-PAGE, and MALDI-TOF-MS (Figure S1). The mutants, Fr-H114A and Fr-S118A, were crystallized under the same conditions used to crystallize FrWT. The X-ray diffraction data, statistics, parameters and RMSDs are summarized in Table S1b.

At –180°C, similar semi-clathrate structures as in FrWT were observed for both Fr-H114A and Fr-S118A (Figure S8). While the semi-clathrate structure of Fr-S118A is essentially identical to that of FrWT (Table S6), Fr-H114A has a structure which more closely resembles that of Maxi than FrWT because an additional water molecule is located near A114 in the space occupied by the imidazole ring of H114. This new water molecule appears to stabilize the semi-clathrate structure by forming hydrogen bonds (Figure S8a). This result indicates that the semi-clathrate (or the pentagonal ring: W1∼W5) is distorted by hydrogen bonding of W3 to the additional water molecule in Fr-H114A. Consequently, the histidine mutation causes shortening of the W2-W3 and W3-W4 distances to 2.36 Å and 2.55 Å, respectively and this causes the pentagonal ring to become less distorted (Table S6). Moreover, in Fr-S118A, the hydrogen bond with the side chain of S118 (anchoring) is lost, but a water molecule is present in the vicinity of the semi-clathrate and forms a hydrogen bond with the semi-clathrate water molecule W4 (Figure S8b). These side chain mutations do not significantly affect the main semi-clathrate structure.

### Dynamic changes of semi-clathrate structures at various temperatures

To evaluate the temperature dependence of the process of formation of the semi-clathrate, the structures of FrWT were determined at various temperatures ranging from -180°C to 20°C (Figures 4 and S10). All structures were refined at 1.6 Å resolution and the refinement statistics are shown in Table S1a. When the temperature was raised to 25°C, we observed minor changes in the cell parameters, indicating expansion of the lattice (Figure S9). A similar lattice expansion was previously reported by Tezcan et al.(34) It was expected that with increasing temperature, loosely bound water molecules would be released or become disordered (i.e., with high B-factors and weak densities), which would ultimately influence the process of formation of the semi-clathrate water structure. The stability of each water molecule was found to vary in a manner depending on its interactions with surrounding residues (side and main chains), the cavity, and other water molecules (Table 1). These factors are reflected in the order of release of semi-clathrate water molecules at high temperatures. We classified the order of water molecule release into four stages with three transition events occurring at -80°C, -40°C and 0°C (Figure 4b, 4d and 4f).

**Figure 4.**
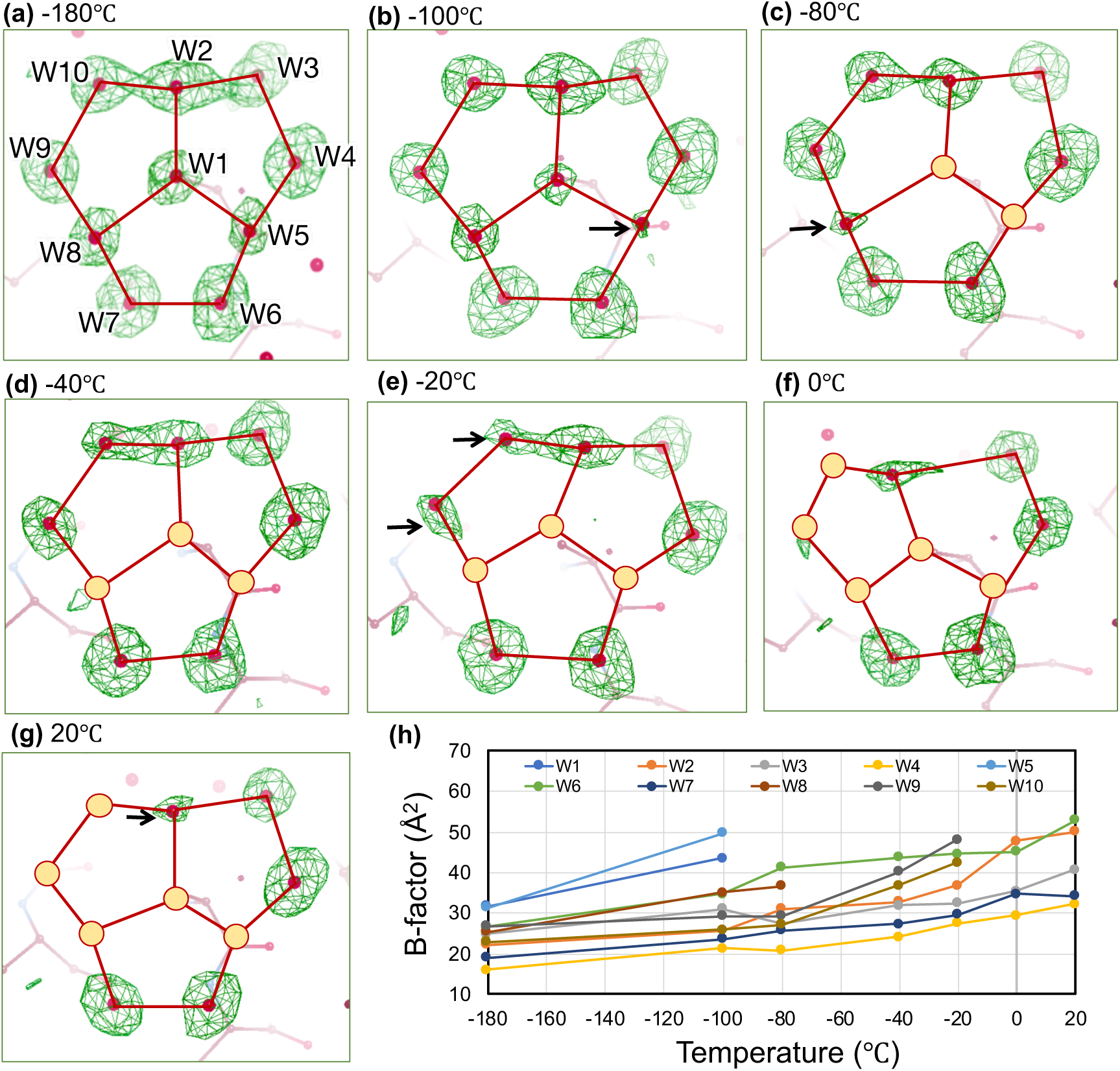
Water release from the semi-clathrate structure of FrWT at various temperatures. (a-g) Structure of semi-clathrate waters in FrWT at -180℃, -100℃, -80℃, -40℃, -20℃, 0℃ and 20℃, respectively. The red circle represents the disappeared water molecule, and the black arrow represents the highly disordered water molecules which are expected to release at next higher temperature. The *F*_o_-*F*_c_ omit density maps are shown in green and contoured at 3.0σ. The images presented here are the screenshots from COOT visualization tool. (h) Changes in the B-factors of the semi-clathrate waters with increasing temperature.

In the first stage (from -180° to -100°C), electron densities attributed to the ten water molecules that formed the semi-clathrate structure were observed (Figure 4a and 4b). The B-factors of the water molecules were found to increase with increasing temperature before release (Figure 4h). At-100°C, the omit map density of W5 was found to decrease significantly (Figure 4b). In addition, the B-factors of W1 and W5 increase significantly from 32 to 44 Å^2^ and 32 to 50 Å^2^, respectively (Figure 4h and Table S3). Such disorder indicates release of these water molecules at higher temperatures. Then, in the second stage (at -80°C), both W1 and W5 disappear while the remaining eight water molecules are maintained in place. W8 becomes disordered at this temperature with a reduced omit density map and an increase in the B-factor from 25 to 37 Å^2^ (Figure 4c and 4h; Table S3). In the subsequent third stage (from -40°C to -20°C), W8 disappears, resulting in a split bistructure (Figure 4d and 4e). As mentioned above, W1, W5 and W8 are weakly stabilized only by three neighboring water molecules within the semi-clathrate (Table 1 and Figure 2b). Due to such low stability, W1, W5 and W8 are released at -40°C. Moreover, in this stage, W9 and W10 are disordered with B-factors of 48 and 42 Å^2^, respectively, indicating partial release (Figure 4e). In the last stage (from 0 to 20°C), W9 and W10 are released. Although W9 is stabilized up to - 40°C by direct hydrogen bonding interactions with Asp112 (Oᵟ), W8 and W10, it becomes more flexible with a high B-factor of 48 Å^2^ at -20°C after the release of W8, prior to its release at 0°C. The remaining five semi-clathrate water molecules are maintained by stabilization from either the main chain, side chains, single anchoring water molecules or the hydrophobic cavity even at 20°C (Table 1). However, among these five water molecules, W2, W3 and W6 each have significantly high B-factors of 50, 53 and 40 Å^2^, respectively, without any direct interaction with residues while W4 and W7 were found to be more tightly fixed in place with relatively low B-factors of 32 Å^2^ and 34 Å^2^, respectively (Figure 4h). W4 and W7 form hydrogen bonds (∼2.8 Å each) with the main chain groups of Ala115 and Asp112, respectively. These hydrogen bonds are similar to those observed at -180°C. Due to such high B-factors, W2, W3 and W6 are expected to be released at slightly higher temperatures. This hypothesis was verified by determining a crystal structure at 25°C which has slightly lower resolution (2.0Å) than others. As a result, W2 and W6 have disappeared and W3 has decreased in density. W4 and W7 remain stabilized by interactions with the main chain (Figure S10).

During temperature variations which cause gradual release of water molecules, the average distances between the remaining semi-clathrate water molecules (W1∼W10) and Ala115 (C^β^) remain consistent with a deviation of less than 0.5 Å (Figure S11 and Table S7). Few changes in side-chain conformations were observed over the temperature variations, while only the B-factors of the water molecules were found to increase before the water molecules are released. These observations indicate that the process of formation of the semi-clathrate structure which occurs with cooling is reversed according to the same order of release of water molecules which occurs with heating. In other words, upon cooling, water molecules such as W3, W4 and W7 that remain until the last stage during heating bind first, followed by other water molecules such as W1, W5 and W8 that dissociate earlier and finally the semi-clathrate structure is completed.

### Reversibility of the process of formation of the semi-clathrate hydrate with cooling and heating

Starting from two different temperatures (−40°C and -20°C) in the third stage, the reversibility in the formation of semi-clathrate hydrate structure was tested, i.e., once cooled to -180°C and then heated to their original temperatures (see Figure 5a where three different positions of a 300-400 µm oil-covered single crystal were used for the X-ray diffraction measurement). At -40°C, three water molecules (W1, W5 and W8) are absent in the initial structure (Figure 5b). This is also observed in Figure 4d. When cooled to -180°C, all of the absent water molecules (W1, W5 and W8) re-appear and form the complete semi-clathrate hydrate structure (Figure 5c) which is similar to the structure shown in Figure 2. In addition, the B-factors of each of the eight semi-clathrate water molecules decrease to less than 30 Å^2^ (except for W5) and are similar to the B-factors of original structure at -180°C as listed in Table S3 (Figure 5e), where W5 tends to dissociate during an early stage (Figure 4b) and thus would have retained a high B-factor of 42 Å^2^. Next, when heated to -40°C, W1, W5 and W8 are absent, as expected (Figure 5d). Anomalies appear in the densities corresponding to W2 and W10 due to a high degree of disorder and therefore only one water molecule is assigned at that location. Moreover, the disappearance and reappearance of water molecules in the semi-clathrate hydrate are reproducible at -20°C and -40°C (Figure 5f-5i). In addition, the slight expansion of lattice parameters is also reversible and varies with temperature (Figure S9). Despite the onset of degradation due to radiation damage and dehydration caused by repeated use of the same crystal in the X-ray diffraction measurements, the resulting crystal structures qualitatively indicate that the process of formation of the semi-clathrate is reversible and temperature dependent.

**Figure 5.**
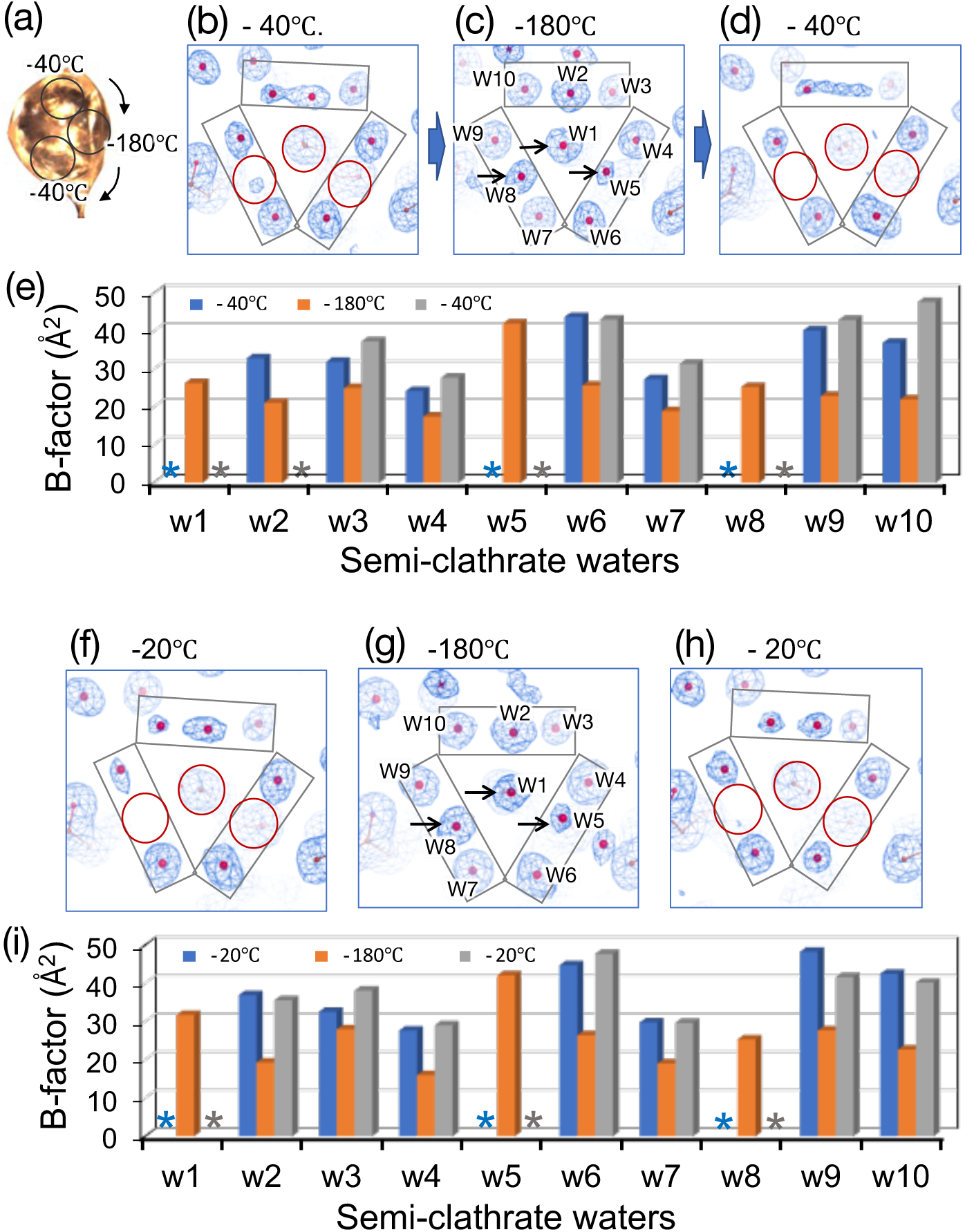
Reversibility in semi-clathrate formation over Ala115. (a) Image of an oil covered FrWT single crystal (∼400µm) used for the variable temperature structure determination. (b-d) Structural changes in the semi-clathrate at -40℃ followed by cooling at -180℃ and further heating to -40℃. (C) Bar diagram showing the changes of B-factors corresponding to the water molecules as described in (b-d). (f) Structural changes in the semi-clathrate over Ala115 at -20℃ followed by cooling to -180℃ and further heating -20℃. (i) Bar diagram showing the changes of B-factors corresponding to the structures in (f-h). The star (*) symbol in the bar diagram represents the disappeared waters. The red circle indicates missing waters. Black arrows represent re-appeared waters after cooling. 2*F*_o_-*F*_c_ maps at 1σ are shown in blue mesh. Images are prepared from COOT.

### Water behaviors observed in molecular dynamics simulations

To verify the order of release of water molecules described above, we conducted molecular dynamics (MD) simulations at different temperatures (from -180 to -30℃) using the -180℃ crystal structure towards better understanding of the semi-clathrate water structure (Figure 6a). The distances between the semi-clathrate water molecules (W1∼W10) from Ala115 (C^β^) were considered as a measure of the stability of the structure. In simulations performed at -180°C and - 130°C, no significant changes in the distances of Ala115(C^β^)-water were observed for 100 ns, indicating minimal changes in the semi-clathrate structure (Figure 6b and 6c). On the other hand, in the simulation performed at -80°C, it was found that W8 first moves away from Ala115 after 5.5 ns, causing the distance between them to reach 12.7 Å at 58.9 ns. This indicates release of W8 at this temperature (Figure 6d and 6e). Moreover, at 100 ns, W1, W5, and W10 also become separated from Ala115 (C^β^) by distances greater than 6-8 Å (Figure 6d) while the other water molecules remain within distances of 5 Å relative to their initial structure. These results agree with information provided by the X-ray structures at -40°C to -20°C in which W1, W5, W8 are completely released and W10 is partially released (with a high B-factor and low density) (Figure 4e). Furthermore, in the simulation at -30°C, it was found that all of the semi-clathrate water molecules are located at distances of >10 Å from the central Ala115 residue, indicating complete disruption of the semi-clathrate structure. This result agrees with information provided by the X-ray structure at 25°C in which only two water molecules (W4 and W7) remain stable, and one water molecule (W3) has decreased density (Figure 6f and S10). The observed discrepancies in water molecule release between the simulations and the experiments may be due to the fact that the crystals were coated with oil to reduce dehydration, and in the simulations, essentially only the solution state of the protein was considered.

**Figure 6.**
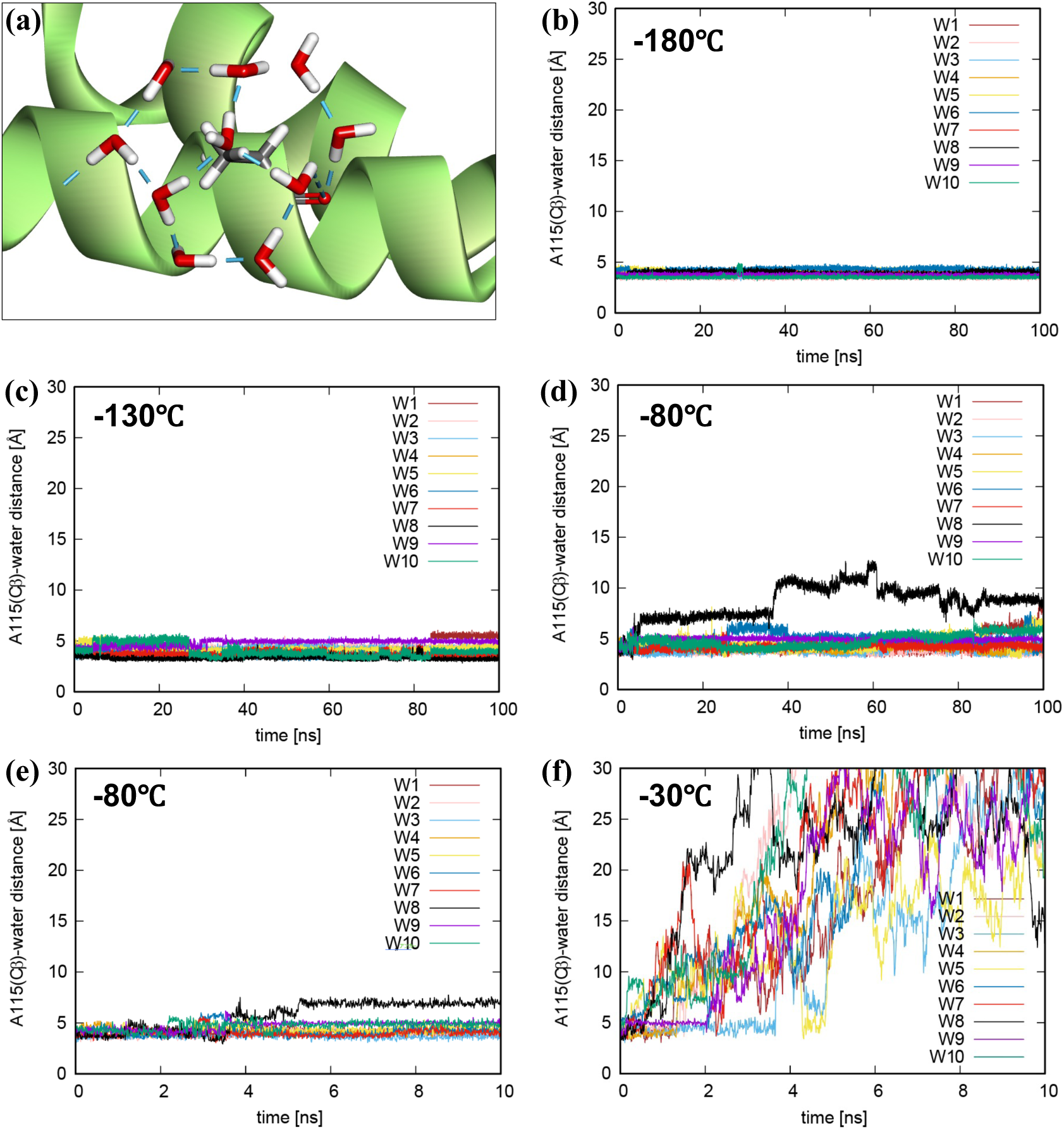
Semi-clathrate water behaviors at various temperatures in MD simulations. (a) Initial structure of semi-clathrate water structure on ferritin cage monomer. (b-d) Time courses of the distances between Ala115(C^β^) and semi-clathrate waters (Ala115(C^β^)-water) at -180℃, -130℃ and -80℃ for 100 ns, respectively. (e-f) Those at -80℃ and -30℃ for 10 ns (short time).

Since the semi-clathrate is located near the interface between the two monomers (Figure 2b), we also investigated the effects at the interface by simulations at -180°C and -80°C with the ferritin cage trimer as the initial structure (Figure S12). In the simulation performed at -180°C, the semi-clathrate water molecules remain quite stable with essentially no change from their initial positions at all sites. On the other hand, in the simulation at -80°C, four water molecules (W5, W6, W8 and W9) at site-2 and six water molecules (W2, W5, W6, W7, W8 and W10) at site-3 move away from Ala115 to distances of 5.8-12.3 Å. In this analysis, we focused only on large changes and noted that essentially all water molecules are maintained at site-1. W2, W6 and W7, remain in position at relatively high temperatures in the experiments but move away from Ala115 in the simulations. The individual dynamics of W5 and W8, (which disappear at -40°C and -80°C, respectively in the experiments), and W9 and W10 (which disappear at 0°C in the experiments) are similar to the results of the simulations. Moreover, W2 and W3, which are located at the interface of the ferritin monomers, remain in position in all the monomer and trimer simulations at temperatures less than -30°C, in good agreement with the experiments. Further investigations are planned because stochastic processes in MD simulations make the process of water release stochastic.

Taken together, the simulations and experiments suggest that W1, W5 and W8 (which are stabilized only by intra-clathrate hydrogen bonds) are the most unstable and W2, W3 (located at the interface), and W9 (direct hydrogen bond with side chain) are relatively stable.

## Discussion

Here, we have elucidated the structure of a single semi-clathrate formed by three fused water rings above Ala115 on the surface of the ferritin cage. The semi-clathrate structure was found to be similar to semi-clathrate structures observed in crambin (a hydrophobic protein) and Maxi (a hyperactive antifreeze protein).(18, 20) The ordered five-membered water array in crambin was formed above the hydrophobic Leu18 residue (C^ᵟ^2) and methylene groups of the Arg17 residue located at the hydrophobic intermolecular cleft in the crystal.(18) In contrast to crambin, ferritin is a four-helix bundled protein and the semi-clathrate is formed above an Ala residue located on a helix surface close to the intramolecular cleft. Therefore, hydrophobic residues on the helix surface near the inter- or intramolecular cleft might be necessary for semi-clathrate formation. This arrangement is similar to the structure observed in the four-helix bundled protein Maxi. Due to its small hydrophobic side chain, the Ala115 residue of ferritin permits exposure of a main chain carbonyl group in an arrangement similar to the configuration seen in Maxi. This exposure induces formation of a semi-clathrate structure which is stabilized by other water molecules and surrounding residues. The mutational analysis (Ala115 to Val, Thr and Gly) revealed that a balance between hydrophobicity and the size of the candidate side chain is necessary to induce formation of the semi-clathrate structure. Mutations of residues surrounding the semi-clathrate have no effect on the structure, indicating an influence of confinement in the structure. However, a hydrophobic residue with a suitable size alone is not sufficient because ferritin includes 14 additional alanine residues that do not form semi-clathrate structures (Figure S2). Ala14 and Ala119 each form only one pentagonal water ring. These results indicate a combination of three necessary elements to form the semi-clathrate structure: (i) a small hydrophobic side chain which provides exposure of the main chain to solvent, (ii) a macrocyclic hydrogen bonding network that forms the framework of the structure, and (iii) amino acid side chains and rigid water molecules in the vicinity to stabilize the structure. These key elements found in our study are in agreement with previous theoretical calculations.(23)

Since the observed wat-wat and wat-C^β^(Ala) distances in the semi-clathrate structures of ferritin and Maxi are similar, it is expected that they are formed via similar mechanisms. We determined the atomic structures at various temperatures using an oil covered single crystal to observe the structural transitions occurring upon dehydration at elevated temperatures. Release of water molecules was observed starting from a threshold temperature of ∼173K (−100°C) and the pentagonal rings begin to break as the temperature is increased. A similar phenomenon was previously observed for water networks in crambin at 200K.(35) The observed threshold temperature (173K) of ferritin is slightly above the glass transition temperature of water (135-140K depending on pressure).(36) We validated the reversibility of water attachment-detachment to the semi-clathrate structure in ferritin by controlling the temperature which indirectly reveals the process of formation of the semi-clathrate structure (Figure 5). When lowering the temperature, the water molecules are expected to participate in formation of the semi-clathrate structure according to their stabilities i.e., the most loosely bound water molecule will join last to complete the structure. Although our experiments were performed using single crystals, MD simulations with monomers and trimers support our interpretation that the semi-clathrate hydrate can form within a certain temperature range in solution and its structural stability is temperature dependent, as we observed experimentally.

Our findings from the basic structural analyses of the semi-clathrate combining both experimental and computational methods reveal critical information about the stability and construction of artificial semi-clathrate structure on a protein surface. The unique water network might be responsible for providing the interaction of ice with ferritin as reported previously.(25, 26) If suitable alanine residues are placed within the confined environment, particularly at the interface of two helices, the resulting engineered protein could form a new semi-clathrate structure. Such studies are in progress. This work is expected to lead to design of new cryomaterials.

In the present study, we determined the structure of a semi-clathrate hydrate present in the solvent channel of an apo-ferritin cage. Such semi-clathrates have been identified as building blocks of several natural antifreeze proteins such as Maxi. It is important to understand the structures of semi-clathrate structures in the interest of constructing and developing artificial cryomaterials. The semi-clathrate hydrate structure on the ferritin cage surface was formed by three fused pentagonal water rings above Ala115. A series of X-ray crystal structures identified two important features: (i) the key elements required to construct the structure and (ii) the dynamics of formation of semi-clathrate structures at various temperatures. A central hydrophobic core (Ala or Val) with macrocyclic hydrogen bonding networks interacting with surrounding residues is required to form such a unique water structure. The semi-clathrate water structure was found to be sensitive to temperature and begins to decompose at temperatures above -100°C. The order of release of water molecules at high temperatures is related to the strength of binding with components of the protein. MD simulations show a similar sequence of release of water molecules. This provides indirect evidence for the formation of semi-clathrate when the temperature gradually is gradually lowered. Therefore, the current work clarifies the characteristics and dynamics of semi-clathrate hydrates. These results will guide the design and construction of new artificial semi-clathrate structures associated with protein surfaces and the development of new cryomaterials for biotechnology applications.

## Materials and Methods

Materials and detailed methods are described in supporting information.

## Author Contributions

BM: X-ray diffraction measurement, structure refinement, data analysis, writing the original manuscript. JT: Protein preparation, crystallization, X-ray diffraction measurement and structure refinement, Data analysis. TF: MD simulations, data analysis, writing the original draft. SA: Structure refinement, data analysis. TU: Supervised the work, data analysis, writing original draft.

## Acknowledgments

This work was supported by “JSPS KAKENHI” (Grant No. 22H00347 to T. Ueno; 22H04744 and 20H05438 to B. Maity) and “Grant-in-Aid for Scientific Research on Innovative Areas ‘Molecular Engines’ (Grant No. JP18H05421 to T. Ueno) from Ministry of Education, Culture, Sports, Science and Technology.

