## Supplemental Information for "Structural insights into a single semi-clathrate hydrate formed in a confined environment of porous protein crystal"

[†] Equal contribution.

* Corresponding authors.

### Experimental

#### Materials and method

All the materials were purchased from commercial sources like Wako, TCI etc. and used without any further purification. The UV-visible absorption spectral measurements were performed using UV-2600PC UV-vis spectrometer (Shimazu).

#### Protein expression and purification

Recombinant L-chain horse spleen apo-ferritin (rHLFr) was used for the study. The mutants were prepared by inverse PCR method (iPCR). The protein was expressed in NovaBlue competent cells (Novagen) transformed with the expression vector pMK2.(1) After expression, the protein was isolated by sonication, heating at 70°C for 15min followed by purification through anion exchange chromatography (Q-Sepharose) and size exclusion chromatography (S-300 Sephacryl). Purity of the 24-mer self-assembled ferritin cage was checked by the Native-PAGE (Figure S1). The mass of the protein monomer was determined by the matrix-assisted laser desorption ionization time-of-flight mass spectrometry, MALDI-TOF-MS (Bruker ultrafle Xtreme) (Figure S1).

#### Protein crystallization

The crystallization of Ferritin and the variants was performed by the hanging drop vapor diffusion method.(2) The drops were prepared by mixing (2μl+2μl) of the purified protein solution (20-30mg $ml^{-1}$) with precipitant solution ((0.5–1.0)M $(NH_4)_2SO_4$, (12.5–20.0)mM $CdSO_4$) and incubated at 20°C. The crystals appeared within a day in most cases. *Caution!!*: $CdSO_4$ is toxic and should be handled with care.

#### X-ray crystal structure determination

The X-ray diffraction data of ferritin crystals were collected in Rigaku XtaLaB Synergy-DW (Cu-kα). Before data collection, crystals were soaked into the precipitant solution (1.0M $(NH_4)_2SO_4$, 20mM $CdSO_4$) containing 25% (v/v) ethylene glycol as cryoprotectant for about 30sec. Automatic data reduction was performed with the CrysAlisPro software (v42). The obtained data were scaled using AIMLESS program in CCP4i2.(3, 4) The pdb model, 1DAT was used as a starting model to determine the phase by MOLREP program.(5) The structure was refined using REFMAC5 (version 5.8.0403) in CCP4i2.(6) The refined model was visualized, inspect and re-builded using COOT

0.9.8.5 EL (CCP4) followed by refinement in REFMAC5.(7) The process was repeated until a suitable model is constructed.

Due to insufficient density, the C-terminal residue Asp174 was left unmodelled in some structures and Lys172 was modelled as Ala in all the structures. The positions of Cd ions were assigned based on the anomalous difference map at 4.0 rmsd. Water molecules in most cases were added automatically in COOT with a cut off residual $F_o$-$F_c$ map at 3.0 rmsd.(8) Weaker density water molecules were added manually. After refinement, the water molecules were inspected manually in coot if they were approximately spherical, have at least some densities in the $2F_o$-$F_c$ map at 1.0 rmsd, no short contacts, approximate tetrahedral geometry etc.(8) The positions of water molecules over the sidechain residue, Ala115 were further inspected by refining the structure at relatively low resolution (2.0Å) with a high I/σ value. The semi-clathrate water molecules were further inspected by omit map (phase bias) and polder map (bulk solvent consideration).(9) Selected semi-clathrate waters (W1, W2, W3, W4, W6, W7, W9 and W10) over Ala115 in FrWT_-180°C were replaced by $Na^+$ ion (one by one) and the refinement resulted increase in the B-factor of the replaced atom but no changes were observed in the difference Fourier map. Validation of $Na^+$ coordination structure with check my metal server (https://cmm.minorlab.org/) suggests uncommon geometry like tetrahedral, square planar or free metal ion with a valance of only 0.4 or less.(10) This imply that $Na^+$ ion might not be positioned in the water network over Ala115.

The final model was validated using wwPDBvalidation server and molprobity before deposition into protein data bank (PDB) server. The accession codes of deposited structures are listed in Table S1.

**Variable temperature X-ray diffraction measurement.**

The X-ray diffraction data were collected at various temperatures starting from at -100°C to 25°C. Temperature of the crystal was controlled with an Oxford Cryosystem low-temperature device. In a typical measurement after achieving the desired set temperature, a suitable sized single crystal in a cryoloop (200-500μm) was mounted to the goniometer, centered, and allowed to stay about 2-3min to achieve the set temperature before data collection. Before mounting, the crystals were first soaked with cryoprotectant (precipitant containing 25% ethylene glycol(v/v)) for about 30sec and then covered up with Santovac®oil (Hampton research) to reduce evaporation. Cryoprotectant was used even for the high temperature data (20°C) to provide a similar experimental condition as measurement in cryotemperature for better comparison among the structures. To minimize radiation damage at high temperature data collection, exposure time, set resolution, number of frames etc. were adjusted (Table S1). For cooling-heating experiment (reversibility) as shown in Figure 5 in the main text, the crystal on the goniometer was allowed to remain as it is during the temperature change and re-centered before data collection. It is to be noted that many times the crystal gave no diffraction or bad resolution at high or variable temperature measurement and the obtained highest resolution datasets are described in this paper.

After measurement, the data were processed as described in section 1.4. Occupancy refinement was not applied during refinement (REFMAC5) to explore the degree of positional disorder (B-factor) of the water molecules at high temperatures as it is associated with how tightly a water molecule is bounded to protein. Water molecules with large deviation of B-factors from the surrounding environment represent highly disordered, loosely bounded, and might have partial occupancy. Thus, appeared close contact waters with high B-factors might not exist simultaneously in a single monomer of the 24-mer ferritin cage. It is to be noted that the electron density maps ($2F_o$-$F_c$ or omit) of some of the semi-clathrate water molecules (W1, W5, W8-10) appeared very weak (close to noise) at high temperatures and still we included them in the model with high B-factors to describe the process of water release because those water densities regularly decreased with increasing temperature starting from -180°C and found reversible as shown in Figure 5 in the main text. Therefore, the assigned waters with weak densities at elevated temperatures might be correct. The final model was validated using wwPDBvalidation server before deposition into protein data bank (PDB) server. The accession codes of deposited structures are listed in Table S1.

**MD simulation**

All the molecular dynamics (MD) simulations were performed using Amber16(11) with the Amber ff14SB force field (protein) and TIP4P-Ew model (water), which has a better temperature dependence than TIP3P model.(12) The crystal structure of FrWT (pdb: 8I6L) obtained in this study was used for the initial structures for monomer and trimer systems in MD simulations. Counter ions were added to preserve the electrostatic neutrality, and the systems were fully solvated by the water box. For the monomer system, first, the energy minimization of 300 steps was carried out with positional restraints on heavy atoms. Then, for the different temperatures at -180, -130, -80 and -30°C, 500 ps equilibration under NVT condition and 500 ps equilibration under NPT condition (1 bar) were conducted, respectively, with the same restraints above. Finally, 100 ns production runs were conducted at each temperature, with the restraints on only main chain atoms, considering the crystal state. For the trimer system, 100 ns MD simulations were performed at -180℃ and -80℃ using the same procedure as the monomer system.

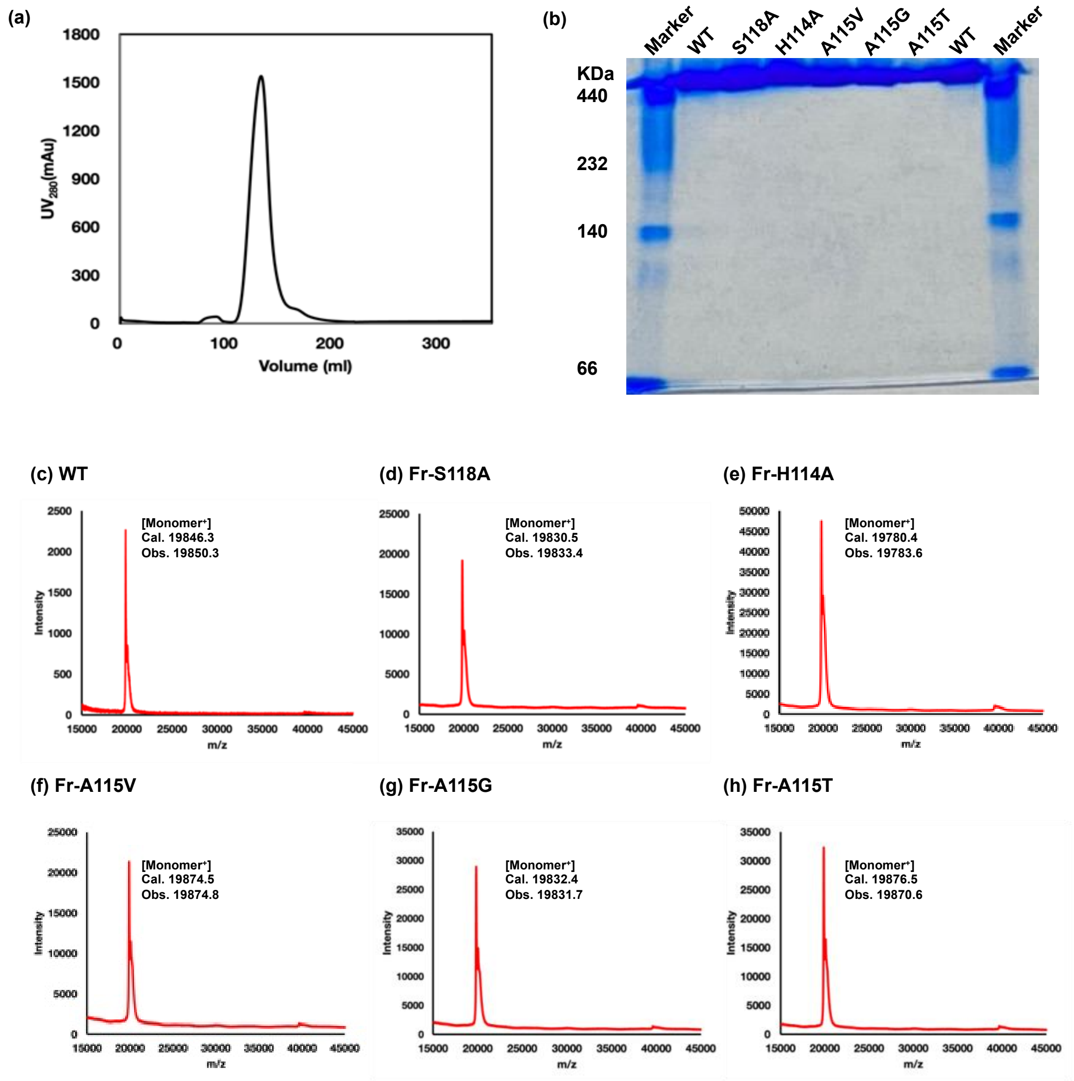


**Figure S1.** Characterization of FrWT and mutants used for the study. (a) A typical elution profile of size exclusion chromatography for FrWT. (b) Native polyacrylamide gel electrophoresis of FrWT and various mutants used for the study. (c-h) MALDI-TOF results for FrWT and various mutants used for the study.

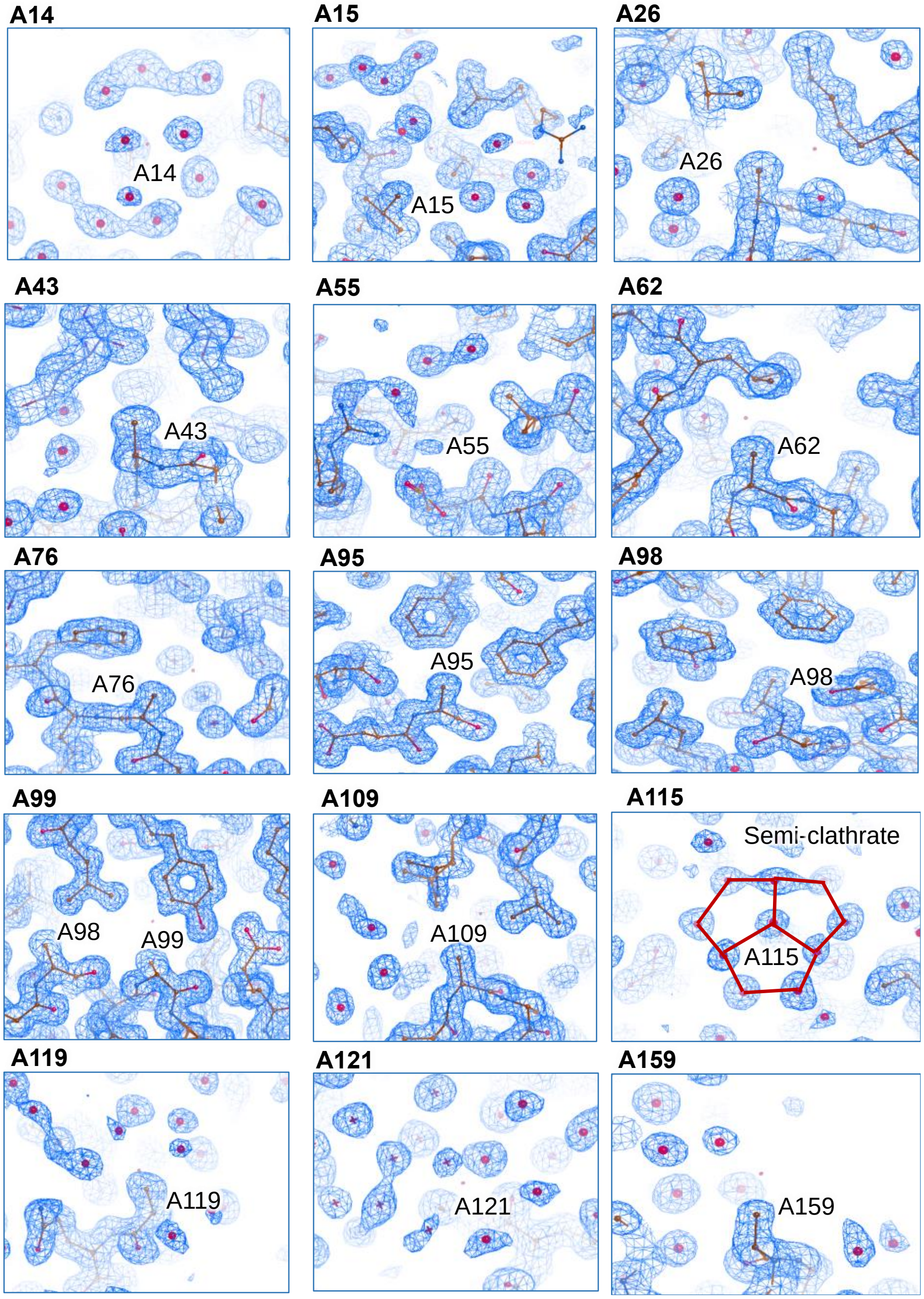


**Figure S2.** Inspection of water molecules surrounding the Alanine residues present in FrWT at -180°C. Out of a total of 15 Ala present in ferritin, only Ala115 was covered with semi-clathrate waters. $2F_o$-$F_c$ maps at 1σ are shown in blue mesh. The solid lines represent the connecting water molecules which forms the pentagonal water rings. The image presented are the screenshots from COOT visualization tool.

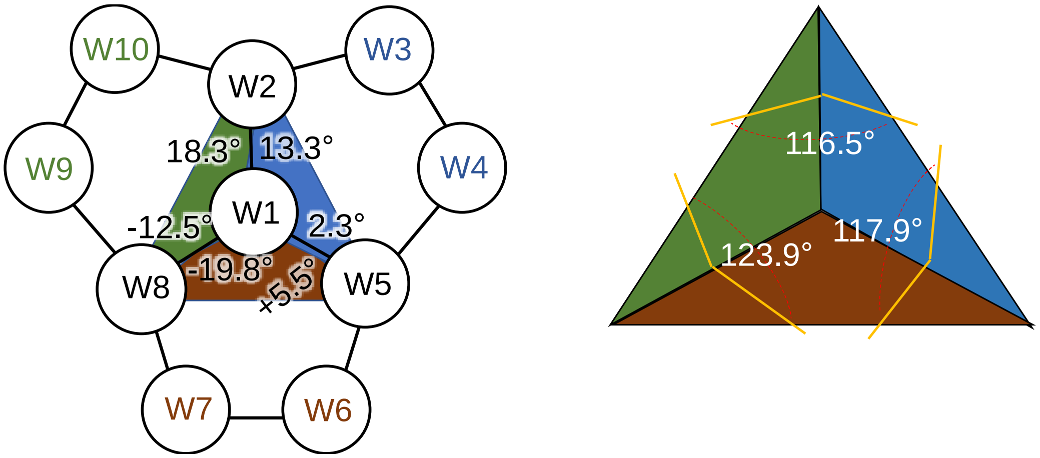


**Figure S3.** Dihedral angles involving four selected water molecules in the pentagonal rings of the semi-clathrate structure of FrWT at -180°C. The selected dihedral angles given in the figure are W8-W1-W2-W10 → 18.3°; W2-W1-W8-W9 → -12.5° W8-W1-W5-W6 → 5.5°; W5-W1-W8-W7 → -19.8° W5-W1-W2-W3 → 13.3°; W2-W1-W5-W4 → 2.3°). Blue, green, and red region are showing the planes consisting of three edge water molecules of the fused section of the polypentagonal rings (W2-W1-W5-W8 → 123.9 °; W8-W1-W5-W2 → 116.5 °; W8-W1-W2-W5 → 117.9 °). The right-side image represents the angles among the pentagonal rings in the semi-clathrate structure. The measurements are done in pymol.

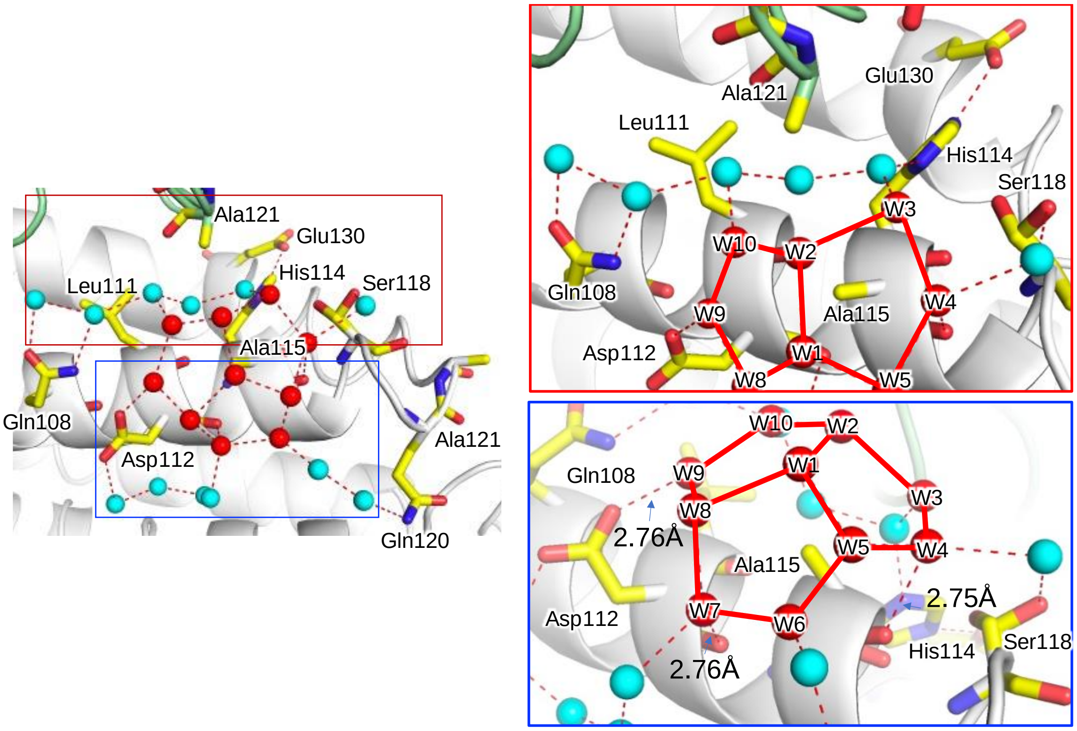


**Figure S4.** Stabilization of the water molecules, W10, W2 and W3 by the hydrogen bonding networks in FrWT at -180°C. The red inset area indicates the stabilization of semi-clathrate W3 by His114 and Glu130. The blue inset area shows the interaction of W4, W6 and W7 through hydrogen bonding interaction with main chain in FrWT. The dashed lines indicate the measured distances using Pymol. The solid lines represent the connecting semi-clathrate waters.

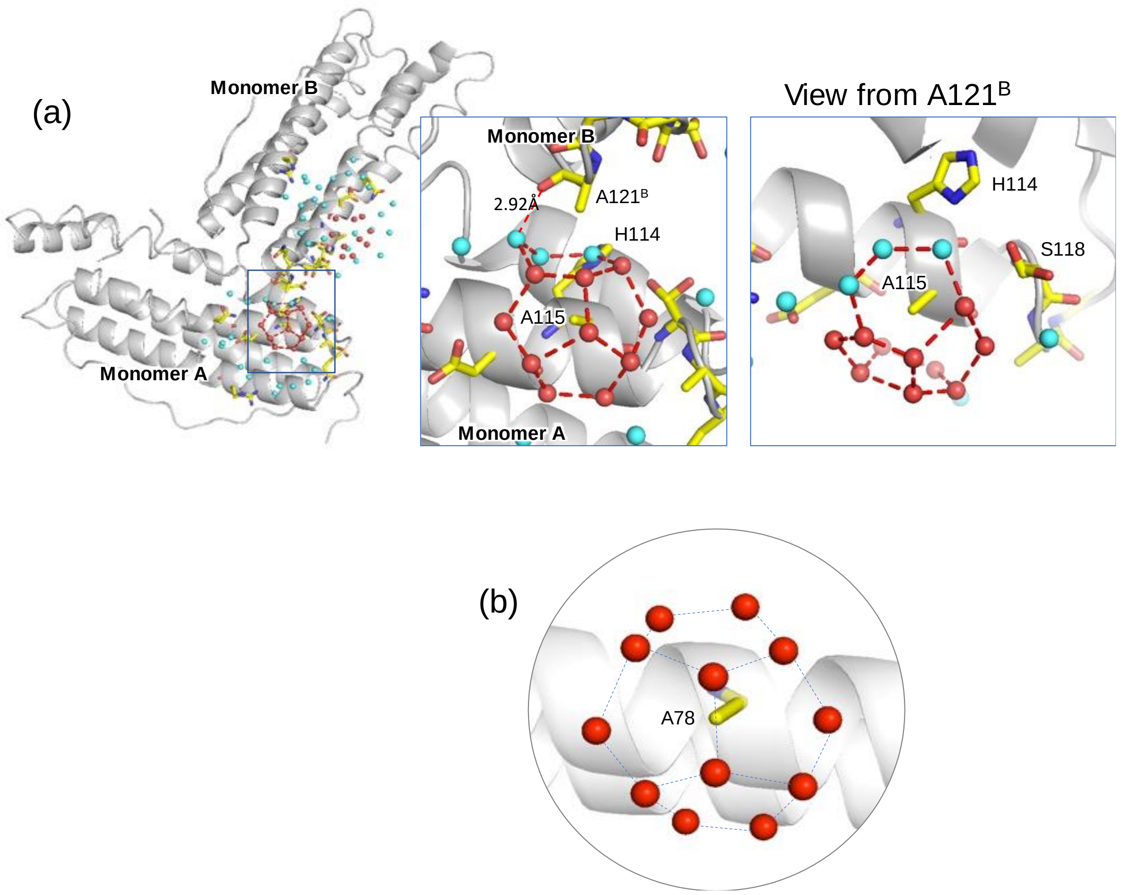


**Figure S5.** (a) Water-water networks near the interface of two ferritin monomers. A unique six-membered water ring was observed at the interface between Ala121 and Ala 115 in FrWT at -180°C. (b) A four fused pentagonal water rings in the semi-clathrate structure over Ala78 in a natural AFP, Maxi (pdb:4ke2).

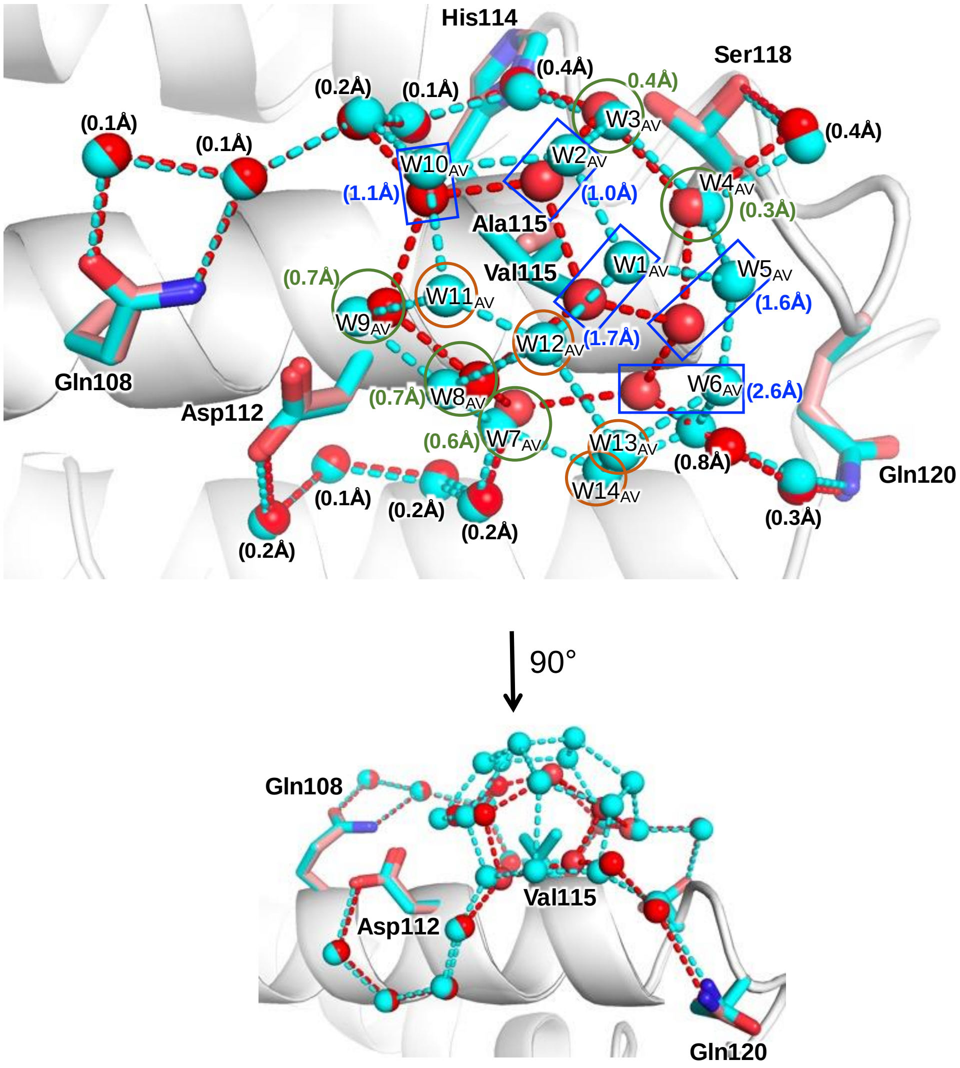


**Figure S6.** Expansion of semi-clathrate water structure in Fr-A115V(Cyan stick/sphere) mutant from FrWT (Red stick/sphere). For simplicity only semi-clathrate waters from Fr-A115V mutant (Cyan spheres) are labelled. The position of ten semi-clathrate water molecules in FrWT are same as Fr-A115V with deviation and marked as green circle and blue rectangle. Green circle and blue rectangle represent the water molecule with small (<1Å) and large movements (>1Å), respectively compared to FrWT in Fr-A115V mutant. The orange circle represents the new water molecules appeared due to A115V mutation. The values given in brackets represent the distance movement from FrWT (red sphere) to Fr-A115V (Cyan sphere). The distances were calculated in pymol after aligning two structures.

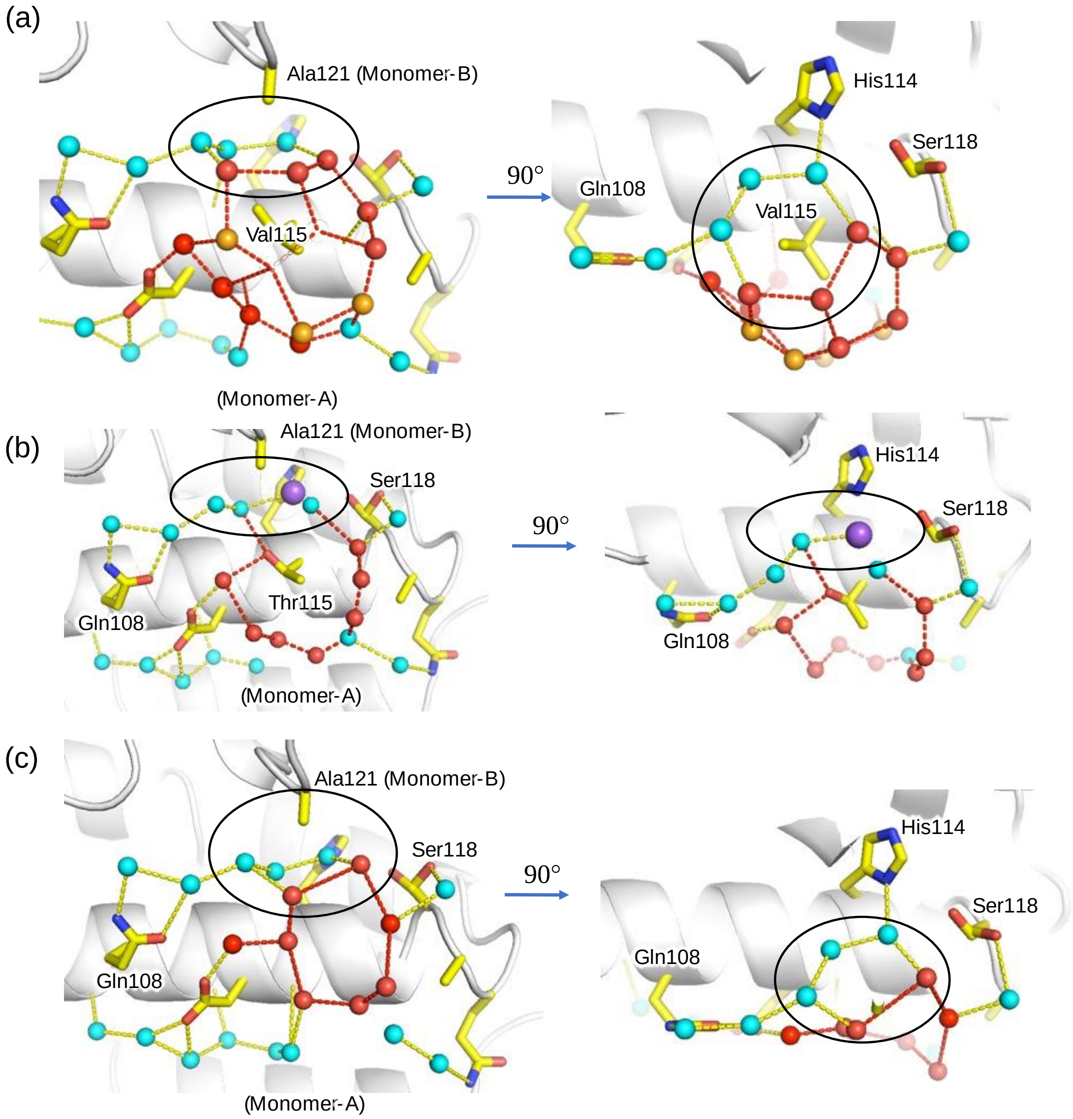


**Figure S7.** Water-water networks (black circle) near the interface of two ferritin monomers. (a) Fr-A115V, (b) Fr-A115T and (c) Fr-A115G.

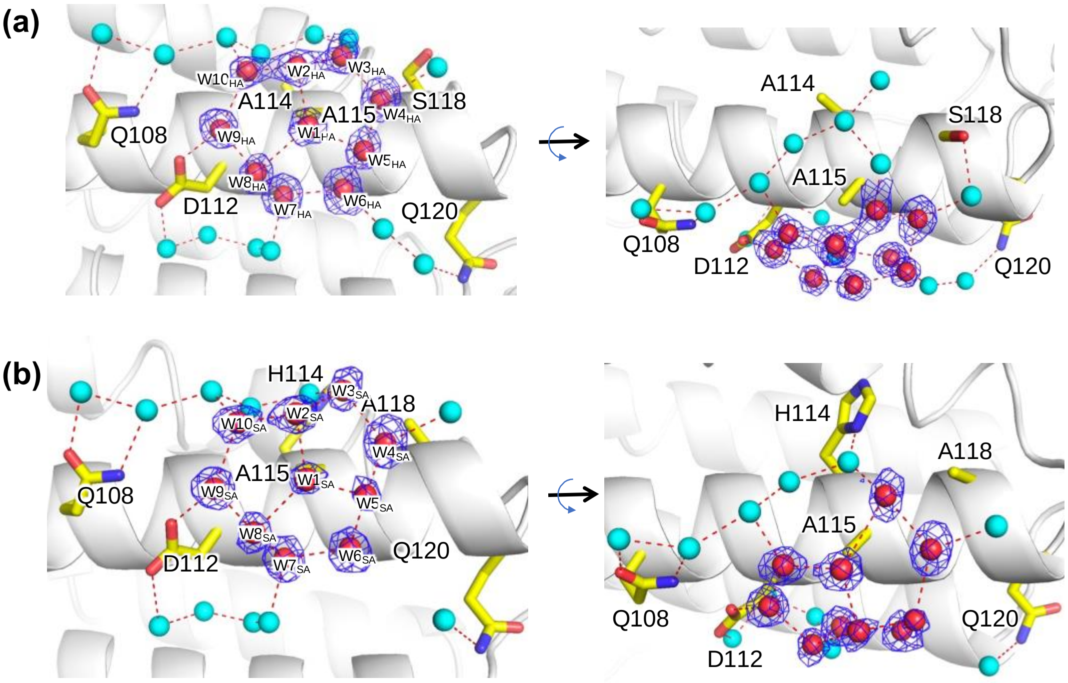


**Figure S8.** Semi-clathrate water structure over Ala115 in (a) Fr-H114A mutant and (b) Fr-S118A mutant. Red sphere indicates semi-clathrate waters and cyan spheres represents anchoring water molecules which stabilizes the semi-clathrate structure. $2F_o$-$F_c$ maps of the selected semi-clathrate waters are shown in blue mesh and contoured at 1σ. The dotted lines represent the connecting waters of interest and having hydrogen bonding distance.

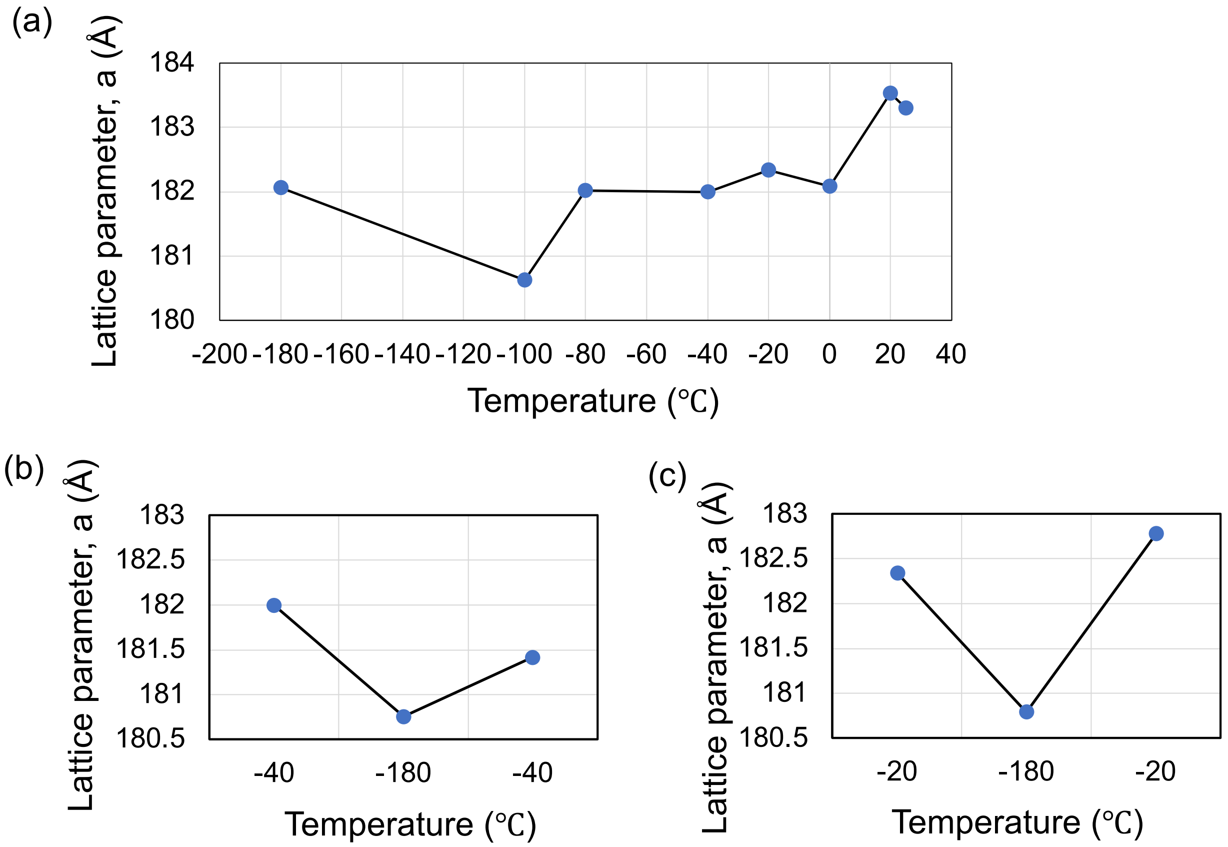


**Figure S9.** (a) Changes of lattice parameters during variable temperature measurements of FrWT crystal as shown in Figure 4a-g. (b-c) Changes of lattice parameters during the reversibility check in the semi-clathrate structure corresponding to the Figure (b-d) and Figure 5(f-h), respectively from the main text. For a cubic system, the lattice parameter, a = b = c.

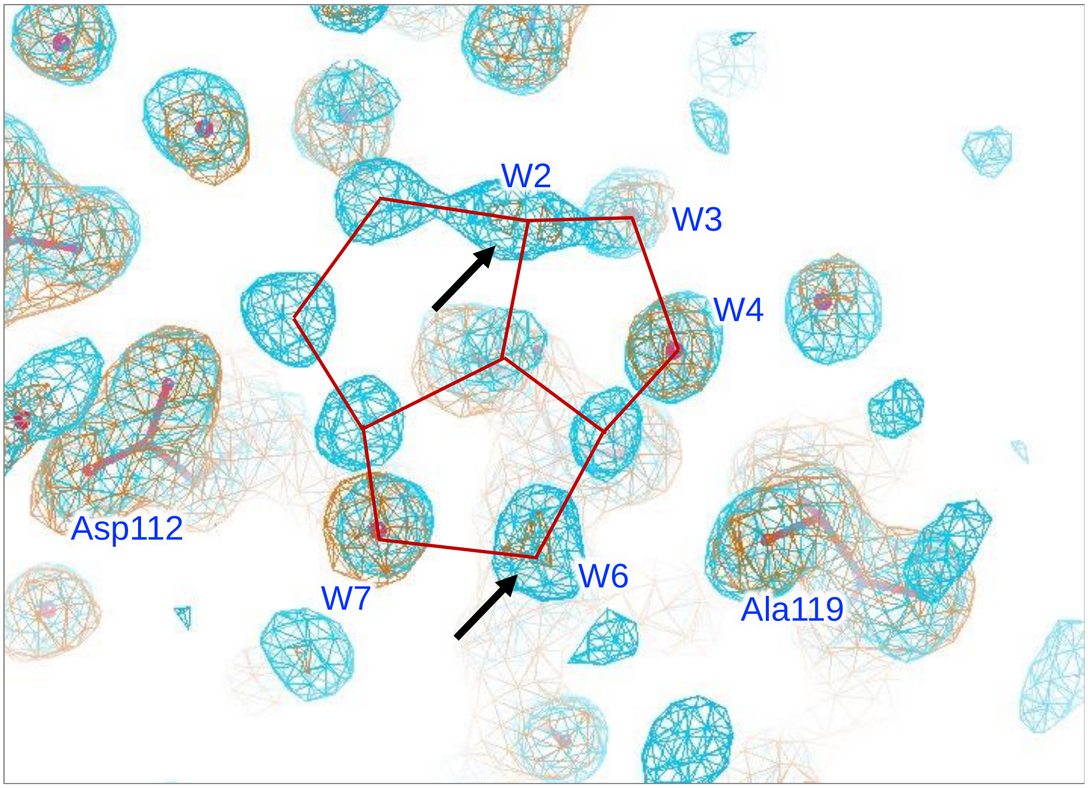


**Figure S10.** Comparison of the water molecule networks surrounding to Ala115 in FrWT at 25°C and -180°C. The structure at 25°C was refined with 2.0Å resolution. The 2$F_o$-$F_c$ map contoured at 1σ for 25°C and -180°C are shown in orange mesh and cyan mesh, respectively. The image was obtained from the COOT visualization tools. The black arrows indicate the release of W2 and W6 at 25°C. The red solid lines represent the semi-clathrate water networks at -180°C.

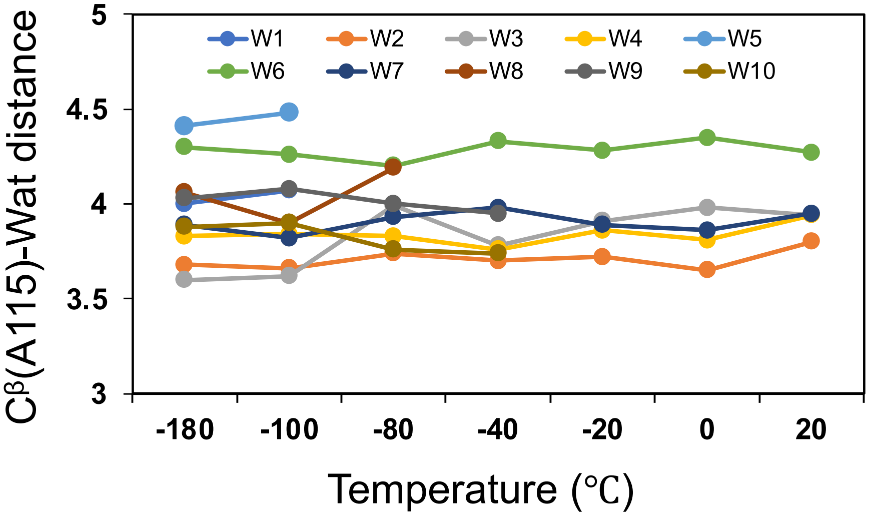


**Figure S11.** Changes in the distances of semi-clathrate waters from $C^{\beta}$ of Ala115 in FrWT with increasing temperatures.

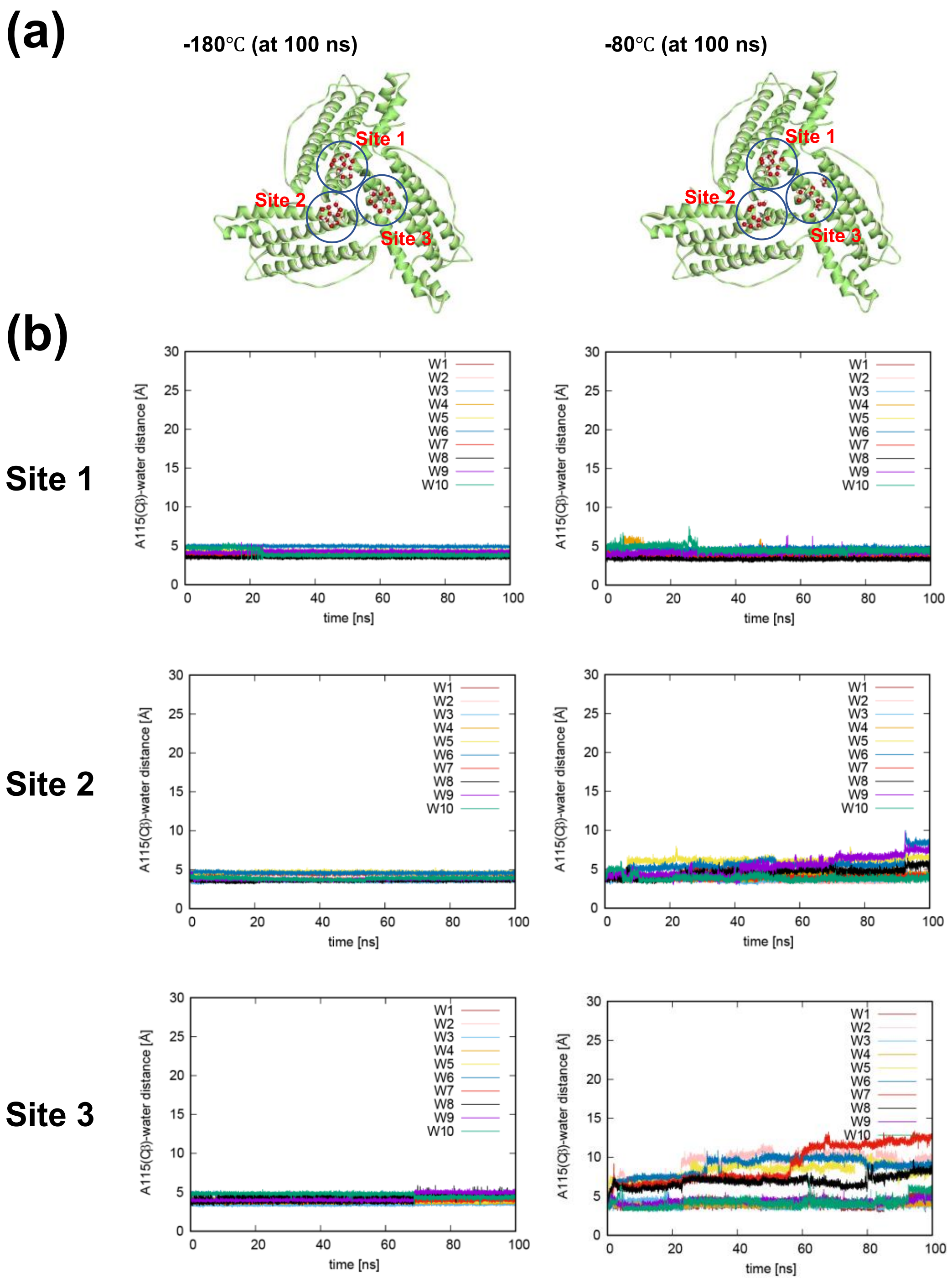


**Figure S12.** MD simulation of the semi-clathrate water structures on ferritin trimer. (a) Snapshot structures of semi-clathrate waters on ferritin trimer at 100 ns in the MD simulations. (b) Time courses of the distances between Ala115($C^{\beta}$) and semi-clathrate water (Ala115($C^{\beta}$)-water) at -180℃ and -80℃ for 100 ns, respectively.

**Table S1a.** Crystallographic data and refinement statics for FrWT at various temperatures

| **Dataset** | **-180°C** | **-100°C** | **-80°C** | **-40°C** | **-20°C** | **0°C** | **20°C** | **-20°C to -180°C (heating)** | **-40°C to -180°C** |
|---|---|---|---|---|---|---|---|---|---|
| **PDB code** | 8I6L | 8J0V | 8J16 | 8J0W | 8J0X | 8J11 | 8J0Y | 8J10 | 8J0Z |
| **Data collection** | | | | | | | | | |
| X-ray source | Cu Kα2 | Cu Kα2 | Cu Kα2 | Cu Kα2 | Cu Kα2 | Cu Kα2 | Cu Kα2 | Cu Kα2 | Cu Kα2 |
| Wavelength (Å) | 1.542 | 1.542 | 1.542 | 1.542 | 1.542 | 1.542 | 1.542 | 1.542 | 1.542 |
| Data collection temperature (°C) | -180 | -100 | -80 | -40 | -20 | 0°C | 20°C | 20°C | 20°C |
| Space group | F432 | F432 | F432 | F432 | F432 | F432 | F432 | F432 | F432 |
| Cell dimensions | | | | | | | | | |
| $a = b = c$ (Å) | 182.058 | 180.624 | 182.015 | 181.999 | 182.340 | 182.084 | 183.529 | 180.786 | 180.76 |
| $\alpha = \beta = \nu$ (°) | 90.00 | 90.00 | 90.00 | 90.00 | 90.0 | 90.0 | 90.0 | 90 | 90 |
| Resolution limit (Å) | 18.58-1.50 (1.53-1.50) | 23.52-1.60 (1.63-1.60) | 23.70-1.60 (1.63-1.60) | 22.23-1.60 (1.63-160) | 22.28-1.60 (1.63-160) | 22.08-1.60 (1.63-1.60) | 22.42-1.60 (1.63-1.60) | 22.09-1.60 (1.63-1.60) | 24.16-1.60 (1.63-1.60) |

| | | | | | | | | | |
|---|---|---|---|---|---|---|---|---|---|
| Unique reflections | 41,630 (1995) | 33,684 (1621) | 34,413 (1651) | 34,339 (1644) | 34,348 (1665) | 34,317 (1619) | 35,274 (1740) | 33,540 (1,633) | 33,140 (1599) |
| Multiplicity | 10.0 (6.7) | 5.2 (3.5) | 4.7 (3.0) | 4.9 (3.0) | 4.7 (3.0) | 4.5 (2.9) | 4.7(3.1) | 4.7 (3.0) | 5.0 (3.3) |
| Completeness (%) | 99.6 (98.2) | 99.7 (98.3) | 99.7 (99.2) | 99.5 (98.0) | 98.9 (97.7) | 99.3 (96.7) | 99.7(97.5) | 99.1 (98.8) | 98.2 (100.0) |
| Mean (I /sigma(I)) | 26.8 (4.1) | 10.6 (1.2) | 10.0 (0.9) | 14.7 (2.5) | 15.6 (1.9) | 13.1 (1.1) | 6.9(0.5) | 10.3 (1.1) | 11.4 (1.6) |
| $R_{meas}$ | 0.057 (0.450) | 0.109 (1.203) | 0.122 (1.461) | 0.069 (0.539) | 0.073 (0.732) | 0.087 (1.102) | 0.132 (2.234) | 0.128 (1.178) | 0.100 (0.766) |
| $R_{merge}$ | 0.054 (0.416) | 0.099 (1.020) | 0.108 (1.195) | 0.062 (0.442) | 0.065 (0.605) | 0.077 (0.901) | 0.1118 (1.850) | 0.114 (0.968) | 0.095 (0.793) |
| $R_{pim}$ | 0.017 (0.170) | 0.046 (0.628) | 0.055 (0.826) | 0.029 (0.303) | 0.031 (0.404) | 0.039 (0.622) | 0.057 (1.232) | 0.057 (0.657) | 0.042 (0.411) |
| Half set correlation CC(1/2) | 1.0 (0.946) | 0.997 (0.508) | 0.993 (0.365) | 0.998 (0.801) | 0.998 (0.708) | 0.997 (0.460) | 0.996 (0.263) | 0.994 (0.521) | 0.997 (0.720) |
| Average mosaicity (°) | 0.78 | 0.64 | 0.61 | 0.77 | 0.70 | 0.69 | 0.67 | 0.71 | 0.73 |

| | | | | | | | | | |
|---|---|---|---|---|---|---|---|---|---|
| Wilson B factor (Å$^2$) | 9.8 | 13.4 | 12.4 | 12.0 | 12.4 | 13.5 | 16.2 | 10.7 | 10.2 |
| **Refinement** | | | | | | | | | |
| Resolution (Å) | 1.50 | 1.60 | 1.60 | 1.60 | 1.60 | 1.60 | 1.60 | 1.60 | 1.60 |
| No. reflections used | 39,495 | 31,986 | 32,664 | 32,577 | 32,627 | 32,573 | 33,451 | 33,845 | 33,503 |
| R-factor/R-free | 0.1549/ 0.1743 | 0.1714/ 0.1905 | 0.1761/ 0.2094 | 0.1453/ 0.1752 | 0.1477/ 0.1687 | 0.1550/ 0.1864 | 0.1686 0.1901 | 0.1797/ 0.2058 | 0.1684/ 0.2015 |
| **No. atoms in the protein** | | | | | | | | | |
| Amino acids | 174 | 173 | 173 | 173 | 173 | 173 | 173 | 173 | 173 |
| Assigned Cd ions | 9 | 3 | 3 | 3 | 3 | 4 | 3 | 3 | 4 |
| Water | 259 | 242 | 233 | 190 | 161 | 142 | 135 | 250 | 254 |
| Ethylene glycol (EDO) | 4 | 4 | 2 | 2 | 1 | 1 | 0 | 4 | 3 |
| **B-factors (Å$^2$)** | | | | | | | | | |
| Overall (protein part) | 11.31 | 15.15 | 14.25 | 13.65 | 14.92 | 16.13 | 19.54 | 12.66 | 11.68 |
| Main chain | 9.69 | 13.10 | 12.27 | 11.43 | 12.37 | 13.22 | 16.16 | 11.01 | 10.20 |

| | | | | | | | | | |
|---|---|---|---|---|---|---|---|---|---|
| Side chains | 12.71 | 17.06 | 16.02 | 15.65 | 17.21 | 18.69 | 22.58 | 14.13 | 12.98 |
| All waters | 25.95 | 30.73 | 30.61 | 29.59 | 29.83 | 32.20 | 35.82 | 25.85 | 25.19 |
| **R.m.s. deviations** | | | | | | | | | |
| Bond lengths (Å) | 0.0131 | 0.0111 | 0.0117 | 0.0111 | 0.0124 | 0.0108 | 0.0101 | 0.0103 | 0.0103 |
| Bond angles (°) | 1.7917 | 1.6221 | 1.5977 | 1.6610 | 1.7140 | 1.6476 | 1.5876 | 1.5632 | 1.6048 |
| **Ramachandran plot statics (%)** | | | | | | | | | |
| Favored region | 98.3 | 98.8 | 98.8 | 98.2 | 98.2 | 98.2 | 98.2 | 98.3 | 98.8 |
| Allowed region | 1.7 | 1.2 | 1.2 | 1.8 | 1.8 | 1.8 | 1.8 | 1.7 | 1.2 |
| Outlier | 0 | 0 | 0 | 0 | 0 | 0 | 0 | 0 | 0 |

Note: Values in the parentheses are for the highest-resolution shell. $R = \Sigma||F_o|-|F_c||/\Sigma|F_o|$, where $F_o$ and $F_c$ are the observed and calculated structure factor amplitudes, respectively. $R_{free}$: and $R$ factor calculated on a partial set that is not used in the refinement of the structure. Ramachandran plot parameters were calculated using RAMPAGE.

**Table S1b**: Crystallographic data and refinement statics for various ferritin mutants.

| **Dataset** | **Fr_A115V** | **Fr_A115T** | **Fr_A115G** | **Fr_S118A** | **Fr_H114A** |
|---|---|---|---|---|---|
| **PDB code** | 8I77 | 8J0U | 8I81 | 8I8U | 8I8Q |
| **Data collection** | | | | | |
| X-ray source | Cu Kα2 | Cu Kα2 | Cu Kα2 | Cu Kα2 | Cu Kα2 |
| Wavelength (Å) | 1.541838 | 1.541838 | 1.541838 | 1.541838 | 1.541838 |
| Data collection temperature (℃) | -180 | -180 | -180 | -180 | -180 |
| Space group | F432 | F432 | F432 | F432 | F432 |
| Cell dimension | | | | | |
| $a = b = c$ (Å) | 181749 | 181.16 | 181.653 | 181.296 | 182.219 |
| $\alpha = \beta = \nu$ (°) | 90.00 | 90.00 | 90.00 | 90.00 | 90.00 |
| Resolution limit (Å) | 30.72-1.50 | 23.58-1.50 | 30.70-1.50 | 18.50-1.6 | 19.10-1.50 |
| | (1.53-1.50) | (1.53-1.50) | (1.53-1.50) | (1.63-1.6) | (1.53-1.50) |

| | | | | | |
|---|---|---|---|---|---|
| Unique reflections | 41,611 (2,029) | 41,193 (2,003) | 41,546 (2,034) | 34,030 (1,652) | 41,908 (2,024) |
| Multiplicity | 7.1 (4.7) | 7.4 (4.8) | 9.6 (6.4) | 4.9 (3.4) | 8.6 (6.3) |
| Completeness (%) | 100.0 (100.0) | 100.0 (99.7) | 100.0 (100.0) | 99.7 (99.7) | 100.0 (100.0) |
| Mean (I /sigma(I)) | 13.8 (1.8) | 19.7 (2.1) | 26.2 (3.6) | 10.8 (1.1) | 25.0 (3.9) |
| $R_{merge}$ | 0.082 (0.742) | 0.075 (0.665) | 0.058 (0.470) | 0.107 (1.072) | 0.064 (0.438) |
| $R_{meas}$ | 0.088 (0.836) | 0.078 (0.748) | 0.061 (0.511) | 0.119 (1.1272) | 0.068 (0.478) |
| $R_{pim}$ | 0.031 (0.380) | 0.028 (0.336) | 0.019 (0.197) | 0.052 (0.671) | 0.022 (0.189) |
| Half-set correlation, CC(1/2) | 0.999 (0.806) | 0.999 (0.742) | 0.999 (0.905) | 0.996 (0.467) | 0.999 (0.918) |
| Average mosaicity (°) | 0.78 | 0.60 | 0.70 | 0.62 | 0.68 |
| Wilson B factor (Å$^2$) | 10.6 | 10.1 | 10.2 | 11.8 | 8.9 |
| **Refinement** | | | | | |
| Resolution (Å) | 1.50 | 1.50 | 1.50 | 1.60 | 1.50 |
| No. reflections used | 39,449 | 39,174 | 39,441 | 39,843 | 39,717 |

| | | | | | |
|---|---|---|---|---|---|
| R-factor/R-free | 0.1601/ 0.1769 | 0.1591/ 0.1808 | 0.1517/ 0.1684 | 0.1710/ 0.2033 | 0.1593/ 0.1741 |
| **No. atoms in the protein** | | | | | |
| Amino acids | 174 | 174 | 174 | 174 | 174 |
| Assigned Cd ions | 5 | 9 | 7 | 4 | 8 |
| Water | 262 | 242 | 253 | 234 | 256 |
| Ethylene glycol (EDO) | 3 | 3 | 4 | 3 | 3 |
| ***B*-factors (Å$^2$)** | | | | | |
| Overall (protein part) | 12.64 | 11.95 | 11.97 | 14.02 | 10.63 |
| Main chain | 11.01 | 10.28 | 10.35 | 12.18 | 8.94 |
| Sidechain | 14.11 | 13.45 | 13.41 | 15.67 | 12.17 |
| Water | 27.27 | 26.67 | 26.57 | 28.15 | 24.39 |
| **R.m.s. deviations** | | | | | |
| Bond lengths (Å) | 0.0120 | 0.0124 | 0.0124 | 0.0107 | 0.0114 |
| Bond angles (°) | 1.6776 | 1.7668 | 1.7057 | 1.5682 | 1.7322 |
| **Ramachandran plot statics (%)** | | | | | |

| | | | | | |
|---|---|---|---|---|---|
| Favored region | 98.8 | 98.8 | 98.8 | 98.8 | 98.3 |
| Allowed region | 1.2 | 1.2 | 1.2 | 1.2 | 1.7 |
| Outlier | 0 | 0 | 0 | 0 | 0 |

Note: Values in the parentheses are for the highest-resolution shell. $R = \Sigma||F_o|-|F_c||/\Sigma|F_o|$, where $F_o$ and $F_c$ are the observed and calculated structure factor amplitudes, respectively. $R_{free}$: an $R$ factor calculated on a partial set that is not used in the refinement of the structure. Ramachandran plot parameters were calculated using RAMPAGE.

**Table S2.** Selected distances of semi-clathrate waters from $C^{\beta}$ of Val115 in FrWT at -180°C.

| | **FrWT** | **Fr-A115V** |
|---|---|---|
| Semi-clathrate water | $C^{\beta}$(Ala115)-Wat dist (Å) | $C^{\beta}$(Val115)-Wat dist (Å) |
| W1 | 4.00 | 5.05 |
| W2 | 3.68 | 4.59 |
| W3 | 3.60 | 4.04 |
| W4 | 3.83 | 4.16 |
| W5 | 4.41 | 5.09 |
| W6 | 4.30 | 5.26 |
| W7 | 3.89 | 3.71 |
| W8 | 4.06 | 3.85 |
| W9 | 4.03 | 4.34 |
| W10 | 3.88 | 4.95 |
| W11 | - | 4.42 |
| W12 | - | 4.96 |
| W13 | - | 5.45 |
| W14 | - | 4.89 |

*Note*: Semi-clathrate in FrWT and Fr-A11V consists of 10 and 14 waters, respectively. W11-W14 are new in Fr-A115V mutant.

**Table S3.** B-factors (Å$^2$) of semi-clathrate waters in FrWT at various temperatures.

| **Semi-clathrate waters** | **B-factors in Å$^2$** | | | | | | |
|---|---|---|---|---|---|---|---|
| | **-180°C** | **-100°C** | **-80°C** | **-40°C** | **-20°C** | **0°C** | **20°C** |
| W1 | 31.76 | 43.60 | - | - | - | - | - |
| W2 | 22.27 | 25.77 | 31.00 | 32.83 | 36.88 | 47.81 | 50.15 |
| W3 | 24.81 | 31.11 | 27.39 | 31.94 | 32.51 | 35.44 | 40.77 |
| W4 | 16.00 | 21.44 | 20.79 | 24.21 | 27.59 | 29.42 | 32.25 |
| W5 | 31.58 | 49.67 | - | - | - | - | - |
| W6 | 26.73 | 34.77 | 41.33 | 43.74 | 44.67 | 45.14 | 52.97 |
| W7 | 19.09 | 23.64 | 25.77 | 27.30 | 29.71 | 34.76 | 34.31 |
| W8 | 25.45 | 35.15 | 36.69 | - | - | - | - |
| W9 | 26.91 | 29.25 | 29.27 | 40.18 | 48.23 | - | - |
| W10 | 22.97 | 25.86 | 27.20 | 36.91 | 42.45 | - | - |

**Table S4.** Selected wat-wat bond distances (Å) in the semi-clathrate water structure of FrWT at variable temperatures and comparison with Maxi.

| **wat-wat interactions** | **FrWT** | | | | | | | **MAXI** |
|---|---|---|---|---|---|---|---|---|
| | **-180°C** | **-100°C** | **-90°C** | **-40°C** | **-20°C** | **0°C** | **20°C** | **-180°C** |
| W1-W2 | 2.72 | 2.89 | - | - | - | - | - | 2.8 |
| W1-W5 | 2.63 | 2.68 | - | - | - | - | - | 3.0 |
| W1-W8 | 3.14 | 3.05 | - | - | - | - | - | 3.1 |
| W2- W3 | 3.28 | 2.94 | 3.82 | 3.47 | 3.54 | 4.35 | 3.95 | 2.6 |
| W3-W4 | 2.93 | 2.74 | 3.07 | 2.95 | 3.03 | 2.94 | 2.97 | 2.7 |
| W4-W5 | 2.91 | 3.05 | - | - | - | - | - | 2.9 |
| W5-W6 | 2.51 | 2.80 | - | - | - | - | - | 2.7 |
| W6- W7 | 2.89 | 2.84 | 3.10 | 2.93 | 3.03 | 2.99 | 3.11 | 2.8 |
| W7-W8 | 2.63 | 2.45 | 2.35 | - | - | - | - | 2.8 |
| W8-W9 | 2.57 | 2.55 | 2.54 | - | - | - | - | 2.8 |
| W9-W10 | 2.77 | 2.87 | 2.67 | 2.75 | 2.68 | - | - | 2.6 |
| W10-W2 | 2.44 | 2.53 | 2.44 | 2.17 | 2.30 | - | - | 2.6 |

Note: Semi-clathrate of MAXI at Ala71 (Figure 1a in the main text) was used for distance measurement.

**Table S5.** B-factors and wat-wat distance in the water networks over Val115 in Fr-A115V mutant.

| **B-factor measurement** | |
|---|---|
| | B-factors (Å$^2$) |
| $W1_{AV}$ | 33.63 |
| $W2_{AV}$ | 21.12 |
| $W3_{AV}$ | 22.62 |
| $W4_{AV}$ | 16.06 |
| $W5_{AV}$ | 25.4 |
| $W6_{AV}$ | 29.87 |
| $W7_{AV}$ | 21.73 |
| $W8_{AV}$ | 35.60 |
| $W9_{AV}$ | 35.99 |
| $W10_{AV}$ | 18.00 |
| $W11_{AV}$ | 39.34 |
| $W12_{AV}$ | 37.25 |
| $W13_{AV}$ | 48.67 |
| $W14_{AV}$ | 28.58 |

| **Distance measurement** | |
|---|---|
| Wat-Wat interaction | Distance in Å |
| $W1_{AV}$ -$W2_{AV}$ | 2.72 |
| $W1_{AV}$ -$W12_{AV}$ | 2.37 |
| $W1_{AV}$ -$W5_{AV}$ | 3.03 |
| $W2_{AV}$ -$W3_{AV}$ | 3.11 |
| $W2_{AV}$ -$W10_{AV}$ | 2.93 |
| $W3_{AV}$ -$W4_{AV}$ | 2.83 |
| $W4_{AV}$ -$W5_{AV}$ | 2.72 |
| $W5_{AV}$ -$W6_{AV}$ | 2.40 |
| $W6_{AV}$ -$W13_{AV}$ | 2.85 |
| $W13_{AV}$ -$W14_{AV}$ | 3.22 |
| $W13_{AV}$ -$W12_{AV}$ | 3.14 |
| $W14_{AV}$ -$W7_{AV}$ | 2.58 |
| $W7_{AV}$ -$W8_{AV}$ | 2.69 |
| $W8_{AV}$ -$W9_{AV}$ | 2.52 |
| $W8_{AV}$ -$W12_{AV}$ | 3.48 |
| $W9_{AV}$ -$W11_{AV}$ | 2.93 |
| $W11_{AV}$ -$W10_{AV}$ | 2.75 |
| $W11_{AV}$ -$W12_{AV}$ | 2.35 |

**Table S6.** Comparison of semi-clathrate wat-wat distance (Å) of FrWT with side chain mutations at residue 114 and 118.

| | **WT-Fr** | **H114A-Fr** | **S118A-Fr** |
|---|---|---|---|
| W1-W2 | 2.72 | 2.73 | 2.82 |
| W1-W5 | 2.63 | 2.73 | 2.54 |
| W1-W8 | 3.14 | 3.26 | 3.18 |
| W2- W3 | 3.28 | 2.36 | 3.56 |
| W3-W4 | 2.93 | 2.55 | 2.98 |
| W4-W5 | 2.91 | 2.84 | 2.98 |
| W5-W6 | 2.51 | 2.47 | 2.53 |
| W6- W7 | 2.89 | 2.79 | 2.84 |
| W7-W8 | 2.63 | 2.52 | 2.45 |
| W8-W9 | 2.57 | 2.58 | 2.53 |
| W9-W10 | 2.77 | 2.83 | 2.81 |
| W10-W2 | 2.44 | 2.39 | 2.48 |

**Table S7.** Changes of semi-clathrate water distance (Å) from $C^{\beta}$ of Ala115.

| | **-180°C** | **-100°C** | **-80°C** | **-40°C** | **-20°C** | **0°C** | **20°C** |
|---|---|---|---|---|---|---|---|
| W1 | 4.00 | 4.07 | - | - | - | - | - |
| W2 | 3.68 | 3.66 | 3.74 | 3.70 | 3.72 | 3.65 | 3.80 |
| W3 | 3.60 | 3.62 | 3.99 | 3.78 | 3.91 | 3.98 | 3.94 |
| W4 | 3.83 | 3.84 | 3.83 | 3.76 | 3.86 | 3.81 | 3.94 |
| W5 | 4.41 | 4.48 | - | - | - | - | - |
| W6 | 4.30 | 4.26 | 4.20 | 4.33 | 4.28 | 4.35 | 4.27 |
| W7 | 3.89 | 3.82 | 3.93 | 3.98 | 3.89 | 3.86 | 3.95 |
| W8 | 4.06 | 3.90 | 4.19 | - | - | - | - |
| W9 | 4.03 | 4.08 | 4.00 | 3.95 | - | - | - |
| W10 | 3.88 | 3.90 | 3.76 | 3.74 | - | - | - |

**Table S8.** Selected distances (Å) in Fr-A115T and Fr-A115G mutants at -180°.

| **Fr-A115T** | | **Fr-A115G** | |
|---|---|---|---|
| $W1_{AT}$-$W2_{AT}$ | 2.56 | $W1_{AG}$-$W2_{AG}$ | 3.23 |
| $W2_{AT}$-$W3_{AT}$ | 2.09 | $W2_{AG}$-$W3_{AG}$ | 3.26 |
| $W3_{AT}$-$W4_{AT}$ | 2.41 | $W4_{AG}$-$W5_{AG}$ | 2.92 |
| $W4_{AT}$-$W5_{AT}$ | 2.18 | $W5_{AG}$-$W6_{AG}$ | 2.34 |
| $W5_{AT}$-$W6_{AT}$ | 2.33 | $W5_{AG}$-$W7_{AG}$ | 2.50 |
| $W6_{AT}$-$W7_{AT}$ | 2.50 | $W6_{AG}$-$W1_{AG}$ | 3.81 |
| $W7_{AT}$-$W8_{AT}$ | 3.16 | $W7_{AG}$-Asp112($O^{\delta}$) | 2.51 |
| $W8_{AT}$-Thr115($O^{\nu}$) | 2.95 | $W4_{AG}$-Asp112 (O) | 2.74 |
| $W8_{AT}$-Asp112($O^{\delta}$) | 2.71 | $W2_{AG}$-Gly115 (O) | 2.68 |
| Na-His114 | 2.91 | | |
| Na- $W9_{AT}$ | 1.74 | | |
